# Metabolic connections between folate and peptidoglycan pathways in *Pseudomonas aeruginosa* inform rational design of a dual-action inhibitor

**DOI:** 10.1101/2023.11.22.568328

**Authors:** Luke N. Yaeger, David Sychantha, Princeton Luong, Shahrokh Shekarriz, Océane Goncalves, Annamaria Dobrin, Michael R. Ranieri, Ryan P. Lamers, Hanjeong Harvey, George C. diCenzo, Michael Surette, Jean-Philippe Côté, Jakob Magolan, Lori L. Burrows

## Abstract

Peptidoglycan is an important bacterial macromolecule that confers cell shape and structural integrity, and a key antibiotic target. The synthesis and turnover of peptidoglycan are carefully coordinated with other cellular processes and pathways. Although there are established connections between peptidoglycan and DNA replication or outer membrane biosynthesis, connections between peptidoglycan and folate metabolism are comparatively unexplored. Folate is an essential cofactor for bacterial growth and required for the synthesis of many important metabolites. Here we show that inhibition of folate synthesis in the important Gram-negative pathogen *Pseudomonas aeruginosa* has downstream effects on peptidoglycan metabolism and integrity. Folate inhibitors reduced expression of the AmpC β-lactamase through perturbation of peptidoglycan recycling, potentiating the activity of β-lactams normally cleaved by that resistance enzyme. Folate inhibitors also synergized with fosfomycin, which inhibits MurA - the first committed step in peptidoglycan synthesis - resulting in dose-dependent formation of round cells that underwent explosive lysis.The insights from this work were used to design a dual-active inhibitor that overcomes NDM-1-mediated meropenem resistance and synergizes with the folate inhibitor, trimethoprim. This work shows that folate and peptidoglycan metabolism are intimately connected and offers new opportunities to exploit this relationship in strategies to overcome antibiotic resistance in Gram-negative pathogens.

## Introduction

High levels of antibiotic resistance in the opportunistic and nosocomial pathogen *Pseudomonas aeruginosa* can limit treatment options and impact the development of new therapeutics^1^. Combination therapy is among the strategies that can be used to restore the efficacy of current antibiotics against resistant strains^2^. For example, β-lactam and β-lactamase inhibitor combinations extended the spectrum of cell-wall targeting β-lactams to strains producing antibiotic-degrading enzymes^3^. Some combinations go beyond unidirectional potentiation, achieving drug-drug synergy through their mutual potentiation. The classic antibiotic combination of trimethoprim (TMP) and sulfamethoxazole (SUL) inhibits separate steps in folate biosynthesis, and together the two drugs are more potent than the single agents^4^. In another example, single-drug inhibition of multiple targets is thought to explain the effectiveness of fluoroquinolones and β-lactams^5^, which essentially synergize with themselves by inhibiting more than one interdependent target.

TMP and SUL block the production of tetrahydrofolate (THF), an essential cofactor in one-carbon metabolism for all forms of life. Humans rely on their diet for folate acquisition, while many bacteria must synthesize folate *de novo* from GTP and chorismate, making the folate biosynthetic pathway an attractive target for antibiotics^6^. Sulfonamide antibiotics, including SUL, inhibit dihydropteroate synthase (FolP) by displacing its substrate, para-aminobenzoic acid (PABA). Some sulfonamides then react with the other FolP substrate, dihydropterin pyrophosphate, to form a dead-end metabolite^7^. FolP is two steps upstream of the TMP target, dihydrofolate reductase (FolA), which catalyzes the formation of THF from dihydrofolate (DHF). THF and its derivatives act as cofactors and one-carbon donors to generate key metabolites including purines, methionine and S-adenosyl methionine, thymidylate, glycine, and serine^8^. The large number of THF-dependent metabolites suggests that treatment with folate inhibitors can have far-reaching effects on cellular physiology. The best-studied effect of folate inhibition is decreased pools of thymidylate, a metabolite required for DNA synthesis^9^. By inhibiting DNA synthesis, TMP treatment can induce the SOS response and expression of SulA, which inhibits assembly of the FtsZ ring that scaffolds the cell division machinery^10^.

A careful review of the literature reveals many clues supporting connections between the folate and PG pathways. Early studies of anti-folates revealed that induction of the SOS-response in folate-depleted cells induced filamentation, suggesting impacts on cell wall metabolism^11,12^. Assembly of the divisome is further regulated by levels of S-adenosyl methionine (SAM), a folate-dependent metabolite^13^. Antifolate-induced morphological changes extend beyond filamentation, as cell-wall deficient cells can also arise following folate inhibition^14^. Further linking the folate and cell wall pathways, Richards and Xing reported accumulation of Lipid II in *Enterobacter cloacae* following TMP and sulfadiazine treatment, suggesting a blockage in PG assembly post-precursor biosynthesis^15^. A screen for *Acinetobacter baumannii* mutants that were significantly less fit during growth with TMP-SUL uncovered mutants with disruptions in peptidoglycan recycling^16^. An attempt to generate antibiotic mechanism-of-action signatures using *B. subtilis* led to misannotation of TMP and SUL as cell wall antibiotics, suggesting their impacts on physiology were similar to those of cell wall-targeting drugs^17^. Synergy between antifolates and cell wall-active antibiotics was reported in multiple studies of drug-drug interactions in *Escherichia coli*, including TMP with vancomycin^18^, oxacillin^19^, fosmidomycin^20^, and mecillinam^21^, and a patent that includes a combination therapy of TMP-SUL and fosfomycin has been issued^22^. Despite these intriguing links, the mechanism(s) of synergy between anti-folates and cell wall-active drugs remains unclear.

As in most Gram-negative bacteria, the peptidoglycan (PG) cell wall of *P. aeruginosa* is composed of repeating disaccharide-pentapeptide subunits that are crosslinked to form a mesh-like structure. PG metabolism can be broadly divided into subunit synthesis, assembly, and turnover/recycling^23–25^. MurA performs the first committed step, transferring an enolpyruvyl group to uridine diphosphate N-acetyl glucosamine (UDP-GlcNAc). MurB-F, MraY, and MurG then catalyze the formation of Lipid II, a GlcNAc-N-acetyl muramic acid (MurNAc) disaccharide with a MurNAc-linked pentapeptide stem (L-alanine [L-Ala], γ-D-glutamate [D-Glu], meso-diaminopimelate [mDAP], and two D-alanine residues [D-Ala]), which is anchored in the inner membrane by undecaprenol diphosphate. MurJ flips Lipid II into the periplasm where transglycosylases attach the GlcNAc of Lipid II to the MurNAc of a growing PG chain. D,D-transpeptidases crosslink PG glycan strands via amide bonds between the 4^th^ (D-ala) and 3^rd^ (mDAP) amino acids of different peptide stems. L,D-transpeptidases crosslink two mDAP residues on different peptide stems, although these 3-3 crosslinks are less abundant than 4-3 crosslinks^26^. Mature PG can be cleaved by amidases, endopeptidases, and lytic transglycosylases to allow for cell growth and division, as well as recycling of PG components. Amidases separate the peptide stem from the glycan backbone, endopeptidases cleave the inter-peptide stem amide bond, and lytic transglycosylases cleave the GlcNAc-MurNAc bond and, for terminal subunits, release a GlcNAc-1,6-anhydroMurNAc (anhMurNAc) peptide. This fragment is taken up, broken down, and its substituents re-enter the PG synthesis pathway. PG metabolism has a high energetic cost, draws upon many different metabolite pools, and carries out the final, irreversible step of the cell cycle; therefore, it is carefully coordinated with many aspects of cell physiology. In this work we use bioinformatics, microscopy, and chemical genetics to characterize the connections between folate and PG metabolism, and leverage our findings in the design of a dual inhibitor that overcomes Gram-negative meropenem resistance.

## Results

We previously discovered that a *P. aeruginosa oprF* mutant was hypersensitive to TMP, suggesting a possible link between folate and PG metabolism^27^. OprF is a major outer membrane porin with a C-terminal PG-binding domain that anchors the outer membrane to the cell wall^28^. The hypersusceptibility of the *oprF* mutant, combined with the results of a detailed literature survey that revealed multiple points of intersection between cell-wall and folate biosynthetic pathways, prompted further investigation. First, we noted several striking structural similarities between a subset of folate and PG enzymes that further strengthen the potential for integration of the pathways. For example, both FolC, the folylpolyglutamate synthase, and MurC-F belong to the Mur ligase family (**Fig 1a**)^29^. MurA is structurally related to AroA (involved in chorismate biosynthesis), and both use the same reaction mechanism for the addition of enolpyruvate^30^. PabC, which catalyzes the production of PABA, is structurally homologous to a D-amino acid aminotransferase^31,32^ that produces D-Glu and D-Ala for the PG stem peptide. Finally, *P. aeruginosa* Cpg2 is a periplasmic carboxypeptidase that cleaves folate to produce glutamyl and pteroate groups^33^; it resembles DapE, which catalyzes production of mDAP^34^. Beyond these structural similarities, there is intriguing synteny between a subset of folate and PG genes (**Fig 1b**). Notably, *folP* is adjacent to *glmM*, encoding a phosphoglucosamine mutase that feeds glucosamine 1-phosphate into PG synthesis. The SUL resistance gene *sul2* (a resistant *folP* allele) is commonly transferred on resistance plasmids with *glmM*^35^.

**Figure 1.**
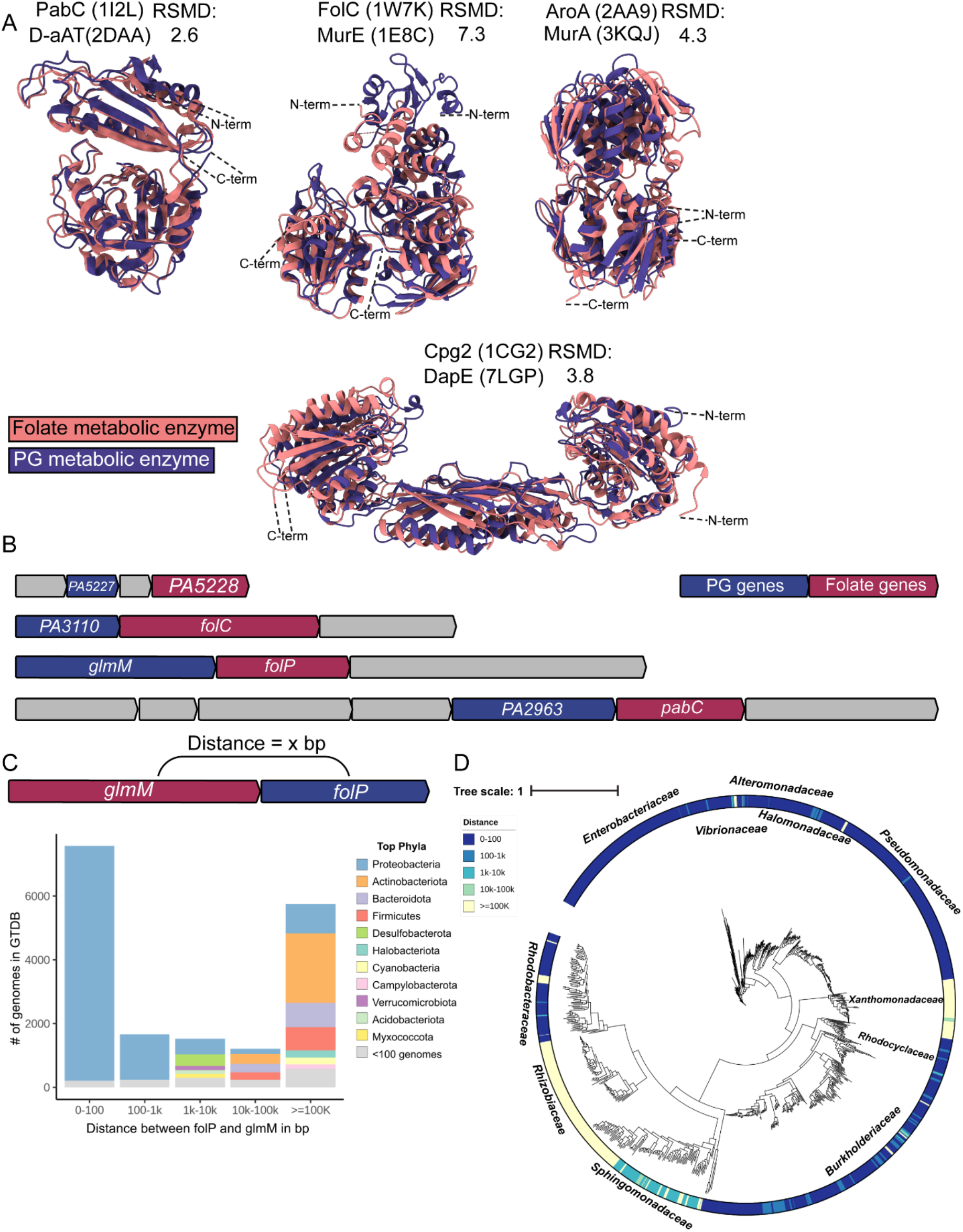
Folate-PG structural and synteny relationships. **A)** A comparison of structural similarity between folate (orange) and PG (blue) metabolic enzymes. The structures were overlayed using the Matchmaker function in ChimeraX. Above each pair of structures are the protein names and PDB codes, where the top name and code corresponds to the folate protein, while the bottom name and code correspond to the PG protein. **B)** An illustration of the *P. aeruginosa* PAO1 operons containing folate and PG-related genes (shown in red and blue, respectively). Other genes within the predicted operons are shown in gray. The unnamed genes are labelled with their *P. aeruginosa* PAO1 locus tag. PA5227 encodes a ZapA homologue, PA5228 encodes a Fau homologue, PA3110 encodes a DedD homologue, and PA2963 encodes an MltG homologue. **C)** The distance in base pairs between the *glmM* and *folP* genes was binned into discrete distances shown on the X-axis, and the number of genomes that fall into each bin are shown on the Y-axis. The legend shows the colour corresponding to each phyla. **D)** An unrooted species phylogenetic tree overlayed with the distance between *folP* and *glmM* mapped across representative genomes of the phylum *Proteobacteria*. The names of major families of the phylum *Proteobacteria* are labelled next to the corresponding genomes. A scale bar and legend are shown in the top left, and the scale bar represents the average number of amino acid substitutions per site. The distance between *folP* and *glmM* is represented by a colour scale, where dark blue indicates a smaller intergenic distance and light yellow indicates a larger intergenic distance.

Synteny can indicate coevolution between genes or pathways^36^. We investigated the extent of the *folP*-*glmM* relationship by measuring the proximity of the two genes across a collection of over 40,000 representative prokaryotic genomes. The results were binned into distances of 0-100, 100-1k, 1k-10k, 10k-100k, and over 100k base pairs. Synteny between *folP* and *glmM* was largely constrained to the phylum *Proteobacteria* (*Pseudomonadota*), but not present in all proteobacteria (**Fig 1c**). Plotting the distance between *folP* and *glmM* on a phylogenetic tree of 938 complete proteobacterial genomes showed that the synteny is generally conserved for all but the families *Xanthomonadaceae*, *Sphingomonadaceae*, and *Rhizobiaceae* (**Fig 1d**). This finding corroborates and expands upon a previous report that *Xanthomonas campestris* is distinct among species of the class *Gammaproteobacteria*, in that *glmM* and *folP* are encoded separately^37^. Some *Rhizobiaceae* species lack a *folP* homologue and presumably import folate, like some lactobacilli^38,39^. The benefits of organizing *glmM* and *folP* together as a transcriptional unit remain unclear, as do the conditions that permit their uncoupling; however, this analysis demonstrates an evolutionary relationship between these genes and prompted additional studies.

### Folate inhibitor phenotypes resemble those caused by cell wall-targeting antibiotics

Following up on our discovery that *a P. aeruginosa oprF* mutant is hypersensitive to TMP^27^, we found that this mutant is also hypersensitive to SUL (**Fig 2a**). We next tested the ability of TMP to compromise envelope integrity using the classic method of exposing cells to hyper- or hypo-osmotic stresses. Changing the NaCl concentration of the LB medium revealed that the minimal inhibitory concentration (MIC) of TMP was ∼8 x lower in high NaCl conditions (**Fig 2b**). To determine if this phenotype was specific to folate inhibition, rather than a general effect of disrupting DNA synthesis, we used the DNA gyrase inhibitor ciprofloxacin (CIP) as a control. The MIC of CIP was reduced ∼2x in response to the same high NaCl stress (**Fig 2c**), suggesting that folate inhibition impacts the cell envelope beyond simply inhibiting DNA synthesis. We reasoned that if TMP exposure affected the cell envelope, it might alter bacterial morphology. PG synthesis inhibitors can have drastic impacts on cell shape and lead to lysis^40–42^. We grew *P. aeruginosa* with increasing concentrations of TMP and examined its morphology using confocal microscopy. TMP induced the formation of round cells that were reminiscent of PG-less L-forms^43^ (**Fig 2d**), suggesting that inhibition of folate synthesis leads to loss of rod shape in a subpopulation of *P. aeruginosa* cells. These round cells eventually undergo explosive cell lysis (**Extended Data Movie 1**). Previous reports of TMP-induced changes in cell morphology suggested increased filamentation; however, that work was primarily done in *E. coli*^42^.

**Figure 2.**
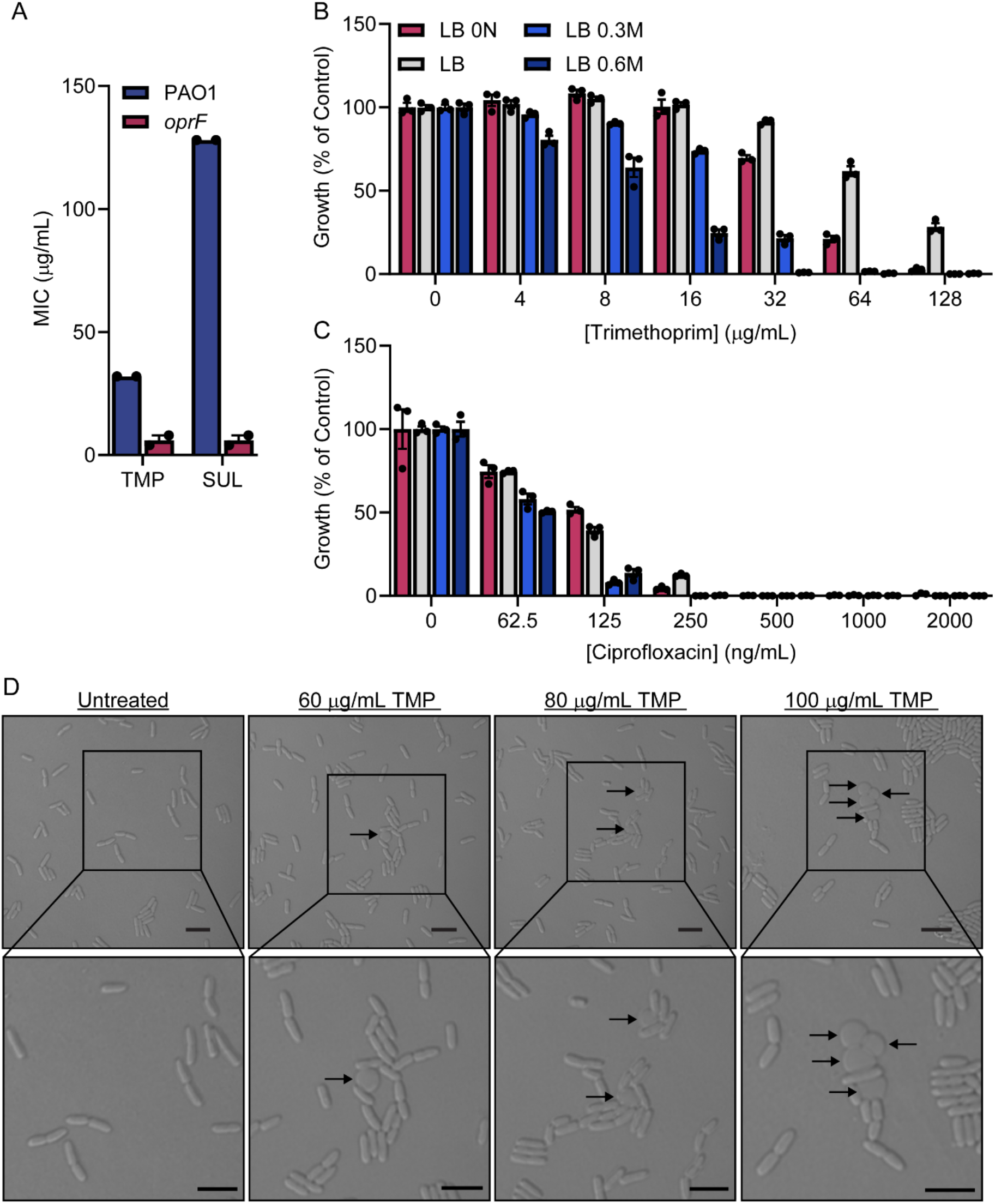
Folate inhibition affects cell envelope integrity. **A)** MICs were measured with a liquid dose-response assay. The antibiotics tested are shown on the X-axis, and the MIC is shown on the Y-axis. The wild type and *oprF* mutant are coloured blue and red, respectively, and individual MICs from two biological replicates are plotted as black circles. The data points for each biological replicate are averaged from three technical replicates. The error bars show the standard error of the mean. **B and C)** A dose-response showing the effect of **B)** TMP or **C)** CIP on PAO1 growth in different osmolarity strength LB. The range of TMP concentrations are shown on the X-axis and growth at each concentration is shown as a percent of the vehicle-treated control and plotted on the Y-axis. Each colour represents different growth media, with the molar concentration of NaCl shown beside the rectangles in the legend (LB 0N = LB with no NaCl). The experiment was repeated in biological duplicates and data from a representative biological replicate are shown, where the bars represent the mean of a technical triplicate, the black circles represent individual data points, and the error bars represent the standard error of the mean. **D)** Micrographs of PAO1 cells treated with increasing TMP concentrations (indicated above each image). A 5-micron scale bar is shown in the bottom right corner of each image. The arrows point to round cells. Close-ups of each image are shown below the original image. Sample preparation and imaging was performed in biological triplicates and images were sampled from at least three separate locations on the agarose pad. Representative micrographs are shown.

### A chemical-genetic screen uncovers specific interactions of PG inhibitors with TMP

Many PG synthesis genes are essential, but sublethal doses of a chemical inhibitor can titrate the activity of these critical enzymes without killing the cell. Therefore, we used a chemical-genetic approach to probe the possible mechanism of antifolate-induced envelope perturbation. Using checkerboard assays, we first screened a collection of PG inhibitors that target different steps of PG metabolism for interactions with TMP. Of the compounds tested (**Fig 3a**), fosfomycin (FOS) and cefoxitin (FOX), which target MurA and multiple penicillin binding proteins (PBPs), respectively (**Fig 3b**), potentiated TMP (**Fig 3c**). This interaction is a general feature of folate inhibition, as SUL also potentiated FOS and FOX (**Fig 3c**). However, FOS and FOX failed to synergize with ciprofloxacin, suggesting that the synergistic effects are not due to inhibition of DNA synthesis (**Extended Data Fig 1a**). Overexpression of FolA, the target of TMP, increased the concentration required for potentiation (**Extended Data Fig 1b**), indicating the primary mechanism of action, rather than off-target effects, drives the interaction. While antifolates and FOS/FOX may synergize by increasing cell permeability (as is the case with potentiation of many antibiotics by polymyxin B), we do not believe this to be the case because: 1) We did not see potentiation of TMP/SUL by polymyxin B (**Extended Data Fig 1c**); 2) Polymyxin B does not potentiate FOS (**Extended Data Fig 1d**); 3) The lack of interaction between TMP and most cell wall-targeting antibiotics (**Fig 3a**) supports more specific mechanisms of potentiation; and 4) no TMP potentiation was observed in a *glpT* mutant that blocks FOS from crossing the inner membrane^44^ (**Extended Data Fig 1e)**.

**Figure 3.**
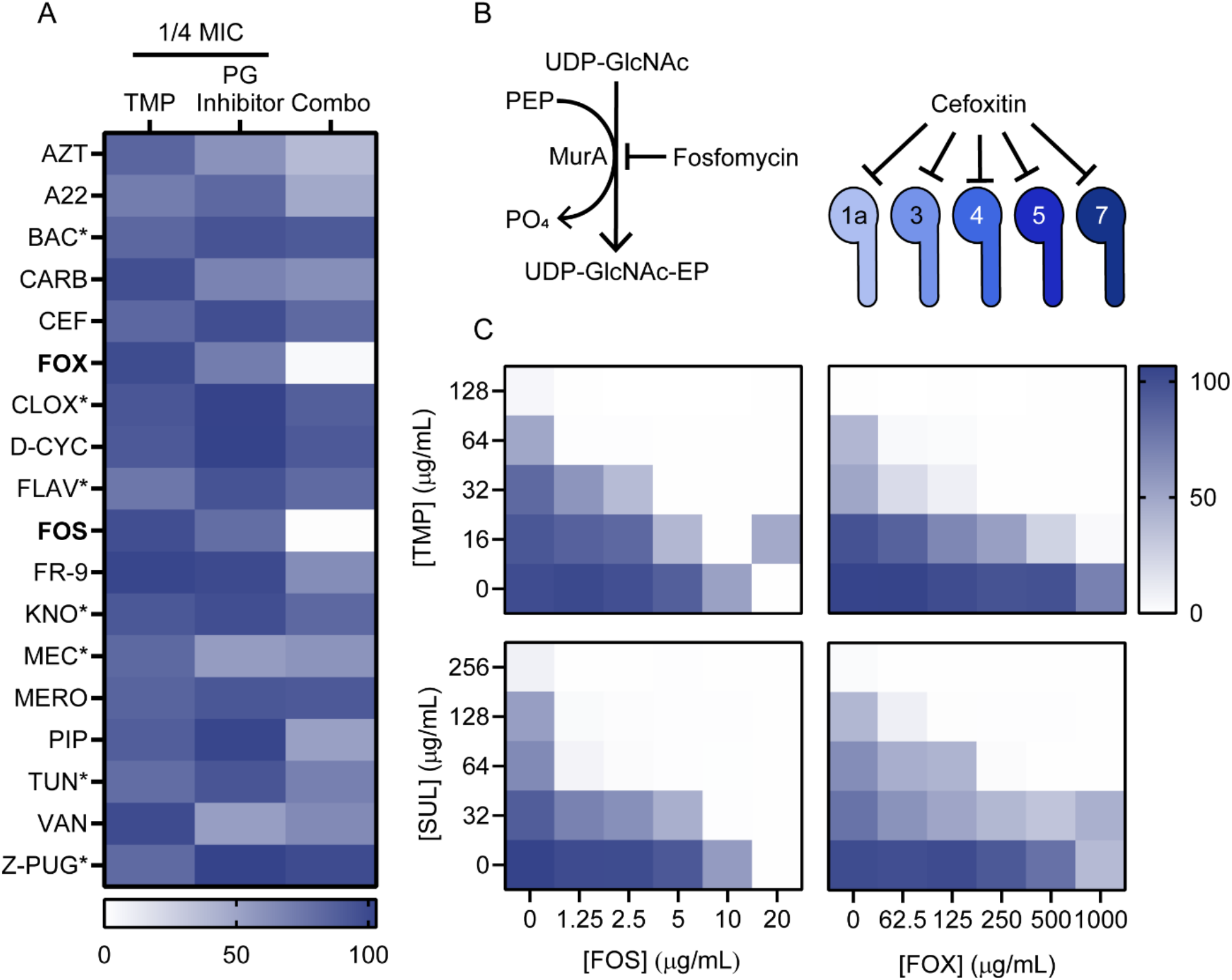
An interaction screen reveals specific antifolate/PG inhibitor potentiation. **A)** The heatmap in **A)** summarizes the data from 8×8 checkerboard assays with TMP and the antibiotics listed to the left of the heatmap. The abbreviations are as follows: AZT=aztreonam, A22=A22, BAC=bacitracin, CARB=carbenicillin, CEF=cefixime, FOX=cefoxitin, CLOX=cloxacillin, D-CYC=D-cycloserine, FLAV=flavomycin, FOS=fosfomycin, FR-9=FR-900098, KNO=kanosamine, MEC=mecillinam, MERO=meropenem, PIP=piperacillin, TUN=tunicamycin, VAN=vancomycin, Z-PUG=Z-PUGNAc. Asterisks indicate antibiotics that did not reach an MIC at the highest concentration tested. Bold font indicates inhibitors that demonstrated synergy. Growth is shown as a percent of the vehicle control and corresponds to the scale bar at the bottom of the heat map. The left column of the heatmap shows growth with ¼ MIC TMP alone. The middle column shows growth after treatment with ¼ MIC of the PG inhibitor alone. The right column shows growth after treatment with ¼ MIC of both TMP and the PG inhibitor. **B)** The left diagram shows the reaction catalyzed by MurA, which is inhibited by fosfomycin. PEP=phosphoenolpyruvate, PO4=phosphate, EP=enolpyruvate. The right diagram shows the PBPs inhibited by cefoxitin, according to Ropy *et al.*^45^. **C)** Heatmaps from 8×8 checkerboards condensed to 5×6 checkerboards. The antibiotics and the respective concentration ranges used are shown on the bottom or left heatmap axes. Growth is shown as a percent of the vehicle control and corresponds to the scale bar on the top right heatmap. Checkerboard assays were repeated in biological triplicates and representative heatmaps from one replicate are shown.

### TMP potentiates FOX by suppressing the AmpR/AmpC response

FOX, like other β-lactams, can inhibit multiple PBPs^45^. We next sought to understand why TMP specifically potentiated FOX, but not the other β-lactams in our panel (**Fig 3a**). *P. aeruginosa* is typically insensitive to FOX, which inhibits PBPs 4 (DacB) and 5 (DacC), D,D-carboxypeptidases that cleave the terminal D-Ala residue from pentapeptide stems to limit the extent of cross-linking. Inhibiting these two PBPs increases the pool of GlcNAc-anhMurNAc-pentapeptides that are recycled via the inner membrane transporter, AmpG. These products bind the regulator AmpR to induce expression of AmpC, a β-lactamase that degrades FOX^46^ (**Fig 4a**). Of the β-lactams we tested, only FOX is both a substrate and inducer of AmpC, suggesting that antifolates might impact the AmpR/AmpC pathway. To test this idea, we chose additional representatives of three types of β-lactams: AmpC inducer <u>and</u> substrate, AmpC non-inducer but substrate, and AmpC inducer but non-substrate. **Figure 4b** shows the fractional inhibitory concentrations (FICI) derived from checkerboard assays of β-lactams from each type in combination with TMP. A pair of compounds with an FICI of 0.5 or less is considered synergistic. Supporting our hypothesis, only those β-lactams that are both inducers <u>and</u> substrates of AmpC synergized with TMP. We reasoned that if AmpC and AmpR were required for the TMP-FOX interaction, then TMP should not further reduce the FOX MIC of *ampR* or *ampC* mutants. Indeed, TMP and FOX failed to synergize in *ampC* or *ampR* mutants (**Fig 4c**).

**Figure 4.**
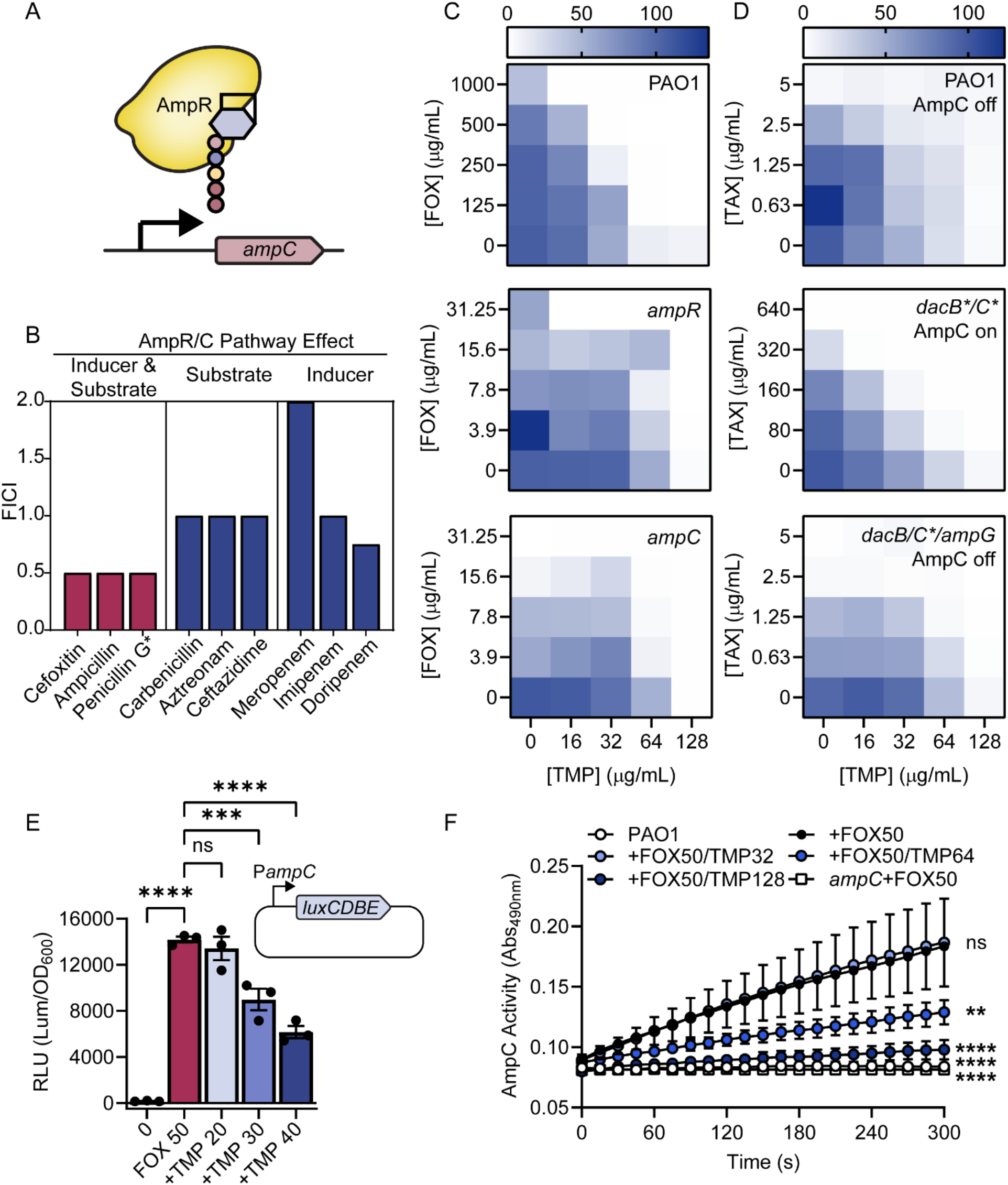
TMP potentiates β-lactams by impairing induction of *ampC* expression. **A)** A schematic showing activation of *ampC* transcription by AmpR bound to anhMurNAc-pentapeptide. **B)** A summary of the fractional inhibitory concentrations (FICI) from 8×8 checkerboard assays of TMP and the antibiotics indicated below each bar. An asterisk indicates that an MIC was not reached for that antibiotic alone. The category of β-lactam is based on whether each β-lactam induces and/or is a substrate for AmpC. The mean FICI was calculated from two biological replicates of 8×8 checkerboard assays. **C)** Heatmaps showing 5×5 checkerboards that condense data from 8×8 checkerboards. The intensity of the blue colour corresponds to the growth as a percent of the vehicle control shown in the legend above the top heatmap. The TMP concentrations are listed at the bottom and FOX concentrations on the left. Note that the FOX concentration range is lower for the *ampC* and *ampR* mutants (middle and bottom) as these mutants are more sensitive. The strain is listed in top right corner of the corresponding heatmap. Checkerboards were performed in biological duplicate, and heatmaps are shown for a representative replicate. **D)** Heatmaps showing 5×5 checkerboards that condense data from 8×8 checkerboards. The intensity of the blue colour corresponds to the growth as a percent of the vehicle control. TMP concentrations are listed at the bottom and TAX concentrations on the left. The TAX concentration range is higher for the *dacBC\*\** mutant as this mutant is more resistant. The strain and status of the AmpC pathway are listed in the top right corner of the corresponding heatmap. Checkerboards were performed in biological duplicate, and heatmaps are shown for a representative replicate. **E)** The level of *ampC* promoter-driven relative luminescence (RLU, calculated as arbitrary luminescence units divided by the matched OD600 growth value) across the different treatment conditions. A schematic showing the plasmid construct containing the *ampC* promoter upstream of the *luxCDBE* genes is shown above the graph. “0” indicates no antibiotic added, FOX50=50 µg/mL cefoxitin, +TMP 20/30/40= 20, 30, or 40 µg/mL trimethoprim added in addition to 50 µg/mL cefoxitin. The bars represent the mean RLU, the individual data points are shown as black circles, and error bars are the standard error of the mean. Technical triplicates were performed for two biological replicates, and a representative replicate is shown. A one-way ANOVA followed by a Dunnett’s multiple comparisons test was performed to compare all the data to the FOX50 condition. ns=not significant, ***=p<0.001, ****=p<0.0001. **F)** Nitrocefin hydrolysis (measured as absorbance at 490 nm) over time. The symbols show the mean of two data points from technical duplicates, the error bars show the standard error of the mean, and the lines connect each symbol over time. The circles show the wild-type strain treated as in the legend. The square symbols indicate the negative control *ampC* treated with cefoxitin. Experiments were performed in technical and biological duplicate, and data from a representative biological replicate are shown.

Genetic inactivation of PBPs 4 and 5 via point mutation of their catalytic Ser to Ala mimics antibiotic inhibition and activates the AmpR/AmpC pathway^45^. Using a PBP4 S72A, PBP5 S64A double mutant (*dacBC*\*\*), we tested whether genetic activation of the AmpR/AmpC response was sufficient for TMP potentiation of cefotaxime (TAX), a non-inducer but substrate of AmpC. As expected, the double mutant was more resistant than the wild type to TAX and we observed TMP potentiation, indicating that inactivating PBP4/5 is sufficient (**Fig 4d**). Further, potentiation was lost in a *dacBC** ampG* triple mutant, suggesting a reliance on AmpG for transport of the inducer GlcNAc-anhMurNAc-pentapeptide into the cytoplasm (**Fig 4d**). A catalytically-inactive S90A point mutant of AmpC also lacked potentiation (**Extended Data Fig 2a**), even though its expression was highly induced by FOX (**Extended Data Fig 2b**), showing that AmpC activity is required for synergy with TMP.

If TMP potentiated FOX activity by preventing *ampC* induction, then transcription from the *ampC* promoter should be reduced. We cloned the *ampC* promoter upstream of a *lux* cassette to create a luminescent transcriptional reporter. As expected, there was almost no luminescence in the untreated condition and very high luminescence in the FOX-treated condition, where *ampC* is induced (**Fig 4e**). Adding TMP in addition to FOX led to a dose-dependent reduction in luminescence. Decreased *ampC* promoter activity implied that AmpC activity in whole cells should also be reduced. We grew cells with FOX alone and in combination with increasing concentrations of TMP, then measured AmpC activity using nitrocefin hydrolysis as a readout. Nitrocefin is a chromogenic β-lactamase substrate commonly used to measure activity^47^. As predicted, we saw a dose-dependent decrease in nitrocefin hydrolysis with TMP relative to the FOX-only control (**Fig 4f**). To rule out the possibility that TMP was a direct inhibitor of AmpC activity, we performed the nitrocefin assay *in vitro* with purified AmpC and TMP, and saw no inhibition (**Extended Data Fig 3**).

### Determinants of the TMP-FOS interaction

With the mechanism of TMP-FOX potentiation clarified, we next addressed the mechanism of TMP-FOS potentiation. Unlike β-lactams, FOS inactivates a single cytoplasmic target (MurA), which can be modelled *in silico*. Flux through folate and PG synthesis pathways can be modelled using genome-scale metabolic reconstruction (GEMs) and predicted with constraint-based flux balance analysis (FBA) using COBRA^48^. We ran an established *P. aeruginosa* GEM through the COBRA FBA to model drug-drug interactions, to determine if TMP/SUL potentiation of FOS could be predicted from known *P. aeruginosa* physiology^49^. The model predicted TMP-SUL potentiation (**Extended Data Fig 4a**), but not the TMP/SUL-FOS interaction (**Extended Data Figs 4b and c**), suggesting that the latter stems from an aspect of cell physiology – possibly PG recycling – not adequately modelled by FBA.

We next looked for genetic interactions with TMP-FOS. First, we found that FOS could not potentiate TMP or SUL against the hypersensitive *oprF* mutant, suggesting that loss of *oprF* and FOS treatment have overlapping effects on antifolate susceptibility (**Extended Data Fig 5**). However, loss of potentiation in the *oprF* background was not particularly informative because OprF has a number of predicted roles^50^. To identify additional mutants with changes in TMP-FOS potentiation, we screened a PA14 transposon mutant library^51^ in 1536-colony density on four agar conditions: no drug, sub-MIC TMP, sub-MIC FOS, or a synergistically lethal combination of the two (**Figs 5a and Extended Data 6a**). This approach allowed us to identify mutants with altered susceptibility to TMP or FOS alone, or mutants such as *oprF* in which TMP-FOS interaction was lost. A subset of mutants grew on the combination plate, suggesting loss of synergy (**Extended Data Fig 6b**). Mutants resistant to only TMP or FOS also grew on the TMP+FOS plates, and we identified those with known changes in susceptibility to TMP or FOS (*anmK*^52^, *oprM*^53^, *folE2*^54^, *glpT*^44^, **Fig 5b**), providing internal validation of the screen. A number of purine biosynthesis mutants, including *purN, purF, purL, purC,* and *purD*, had increased sensitivity to the TMP/FOS combination. We cherry-picked eight purine biosynthesis mutants, including the five above, from the transposon library and performed follow-up checkerboards. Some mutants displayed a small increase in TMP sensitivity, which was expected given that purine biosynthesis depends on folate (**Extended Data Fig 7**). Notably, *purC* and *purM* mutants were 8 and 4 times more sensitive to TMP, while the *purM* mutant was ∼4 times more resistant to FOS. As well, *purD* and *purN* mutants were slightly more sensitive to the TMP/FOS combination. Although it is still unclear how disruption of purine biosynthesis impacts FOS susceptibility, these data suggest a possible mechanism underlying the TMP-FOS interaction.

**Figure 5.**
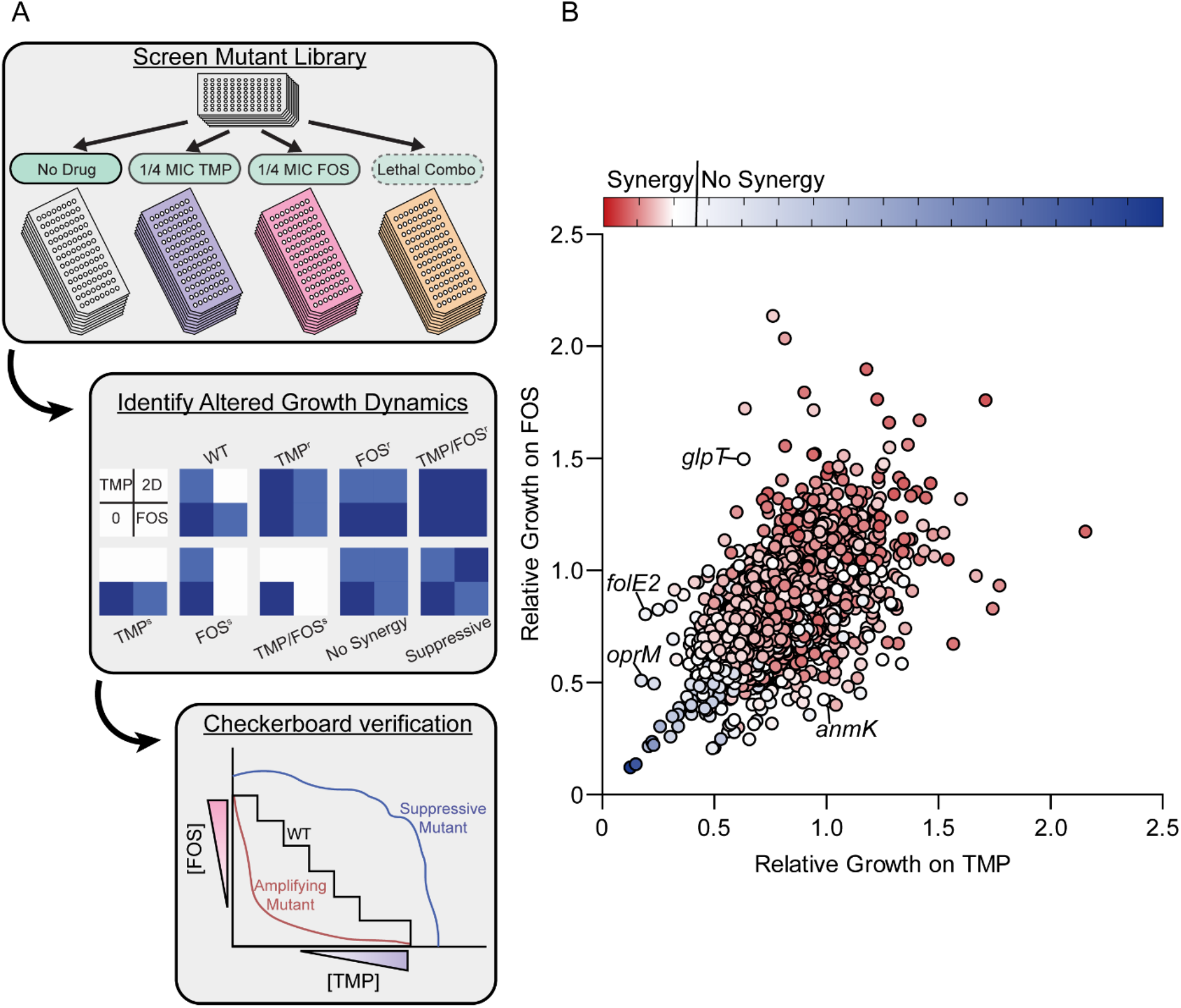
A genome-wide screen of TMP-FOS interaction determinants. **A)** A schematic outlining the mutant library screen, analysis, and follow-up workflow. The top panel shows the four solid medium conditions on which the library was pinned. The middle panel shows the predicted outcomes of mutants in the screen. “2D” indicates the two-drug treated condition, while “s” and “r” indicate hypersensitive or resistant, respectively. Blue represents growth. The bottom panel shows the predicted outcomes for the wild-type strain and different mutants on a checkerboard assay. **B)** A scatter plot showing the results of the PA14 library screen. Normalized growth relative to the untreated control for the TMP (X-axis) or FOS (Y-axis)-only conditions are plotted on the axes. Red and blue indicate synergy or no synergy, respectively, while colour intensity indicates the degree of synergy. Data points for select mutants that internally validate the screen are labelled.

### Disruption of PG recycling by TMP

Aberrant PG recycling is a common determinant of sensitivity to both FOS and FOX in *P. aeruginosa*^55^, consistent with our observation that TMP effects on AmpC expression were AmpG-dependent. To explore this further, we reasoned that treatment with TMP might change the abundance or proportions of soluble PG recycling fragments, and used LC-MS to measure abundance of select soluble PG species. In TMP-treated samples, we observed significantly increased GlcNAc-anhMurNAc and decreased anhMurNAc abundance (**Figure 6)**, suggesting a possible block in PG recycling. Disaccharides lacking the stem peptide are products of amidases, either periplasmic or cytoplasmic, and a substrate for the NagZ β-*N*-acetylglucosaminidase. AmpD amidase activity is negatively correlated with AmpC induction^56^, while NagZ activity is required for FOS resistance^57^, so accumulation of this disaccharide is consistent with known factors that increase FOX and FOS sensitivity.

**Figure 6.**
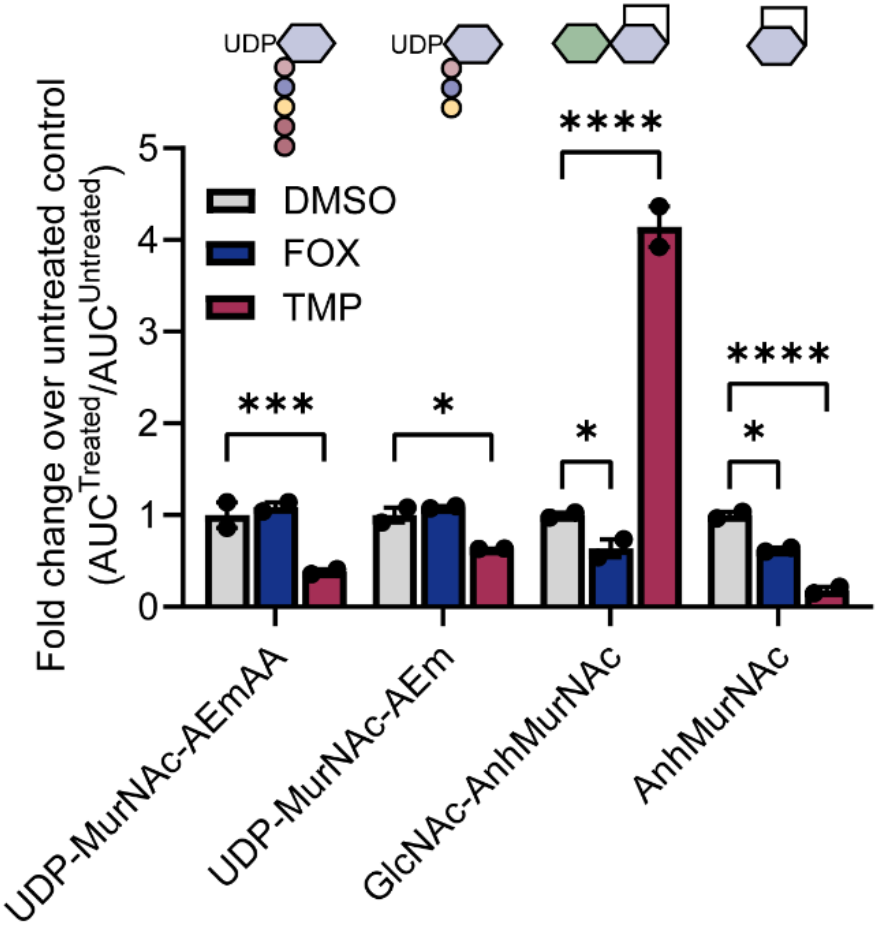
TMP treatment drives accumulation of the GlcNAc-anhMurNAc PG recycling intermediate. The abundance of different soluble PG metabolites was measured by LC-MS and quantified by integrating the peak of the extracted ion chromatogram corresponding to each species’ M/Z. The PG metabolites are listed on the X-axis and a cartoon of each is shown above the corresponding bar. AEmAA = the five amino acids of the stem peptide. The Y-axis shows the integrated peak value or area under the curve (AUC) for each condition and species divided by AUC of the DMSO control sample for the matched species, and the data are represented as the fold-change relative to the control. Therefore, each DMSO condition has a mean fold change of 1. Two biological replicates were performed, each with two technical replicates. One representative biological replicate is shown, and each individual data point is shown as a black circle. The bars indicate the mean of the technical replicates, and the error bars show the standard of the mean. The grey, blue, and red bars correspond to DMSO, FOX (50 µg/mL), and TMP (64 µg/mL) treated samples, respectively. A two-way ANOVA followed by Dunnett’s multiple comparisons test was performed to compare the FOX and TMP treated conditions to the DMSO control. *=p<0.05, ***=p<0.001, ****=p<0.0001.

### A novel dual inhibitor overcomes meropenem resistance by targeting FolP and NDM-1

Having established at least two ways that anti-folates may impact susceptibility to cell wall-targeting antibiotics, we leveraged the folate-PG relationship for rational design of a novel dual-function small molecule inhibitor. The recently developed metallo-β-lactamase (MBL) inhibitor ANT-2681 has some structural resemblance to sulfathiazole^58^, a sulfonamide antibiotic that inhibits folate synthesis (**Fig 7a**)^59^. ANT-2681 contains the core 2-sulfanilamidothiazole of sulfathiazole with some additional substituents, including a carboxylate moiety that is required for MBL inhibition via coordination to active-site zinc ions. The key structural feature of sulfathiazole is the sulfanilamide that competes with *para*-aminobenzoic acid (PABA) for the FolP-binding pocket^59^. An X-ray crystal structure of sulfathiazole-bound FolP showed that the variable ring of sulfonamides sits outside the binding pocket and thus could likely tolerate substitution on the thiazole moiety^59^.

**Figure 7.**
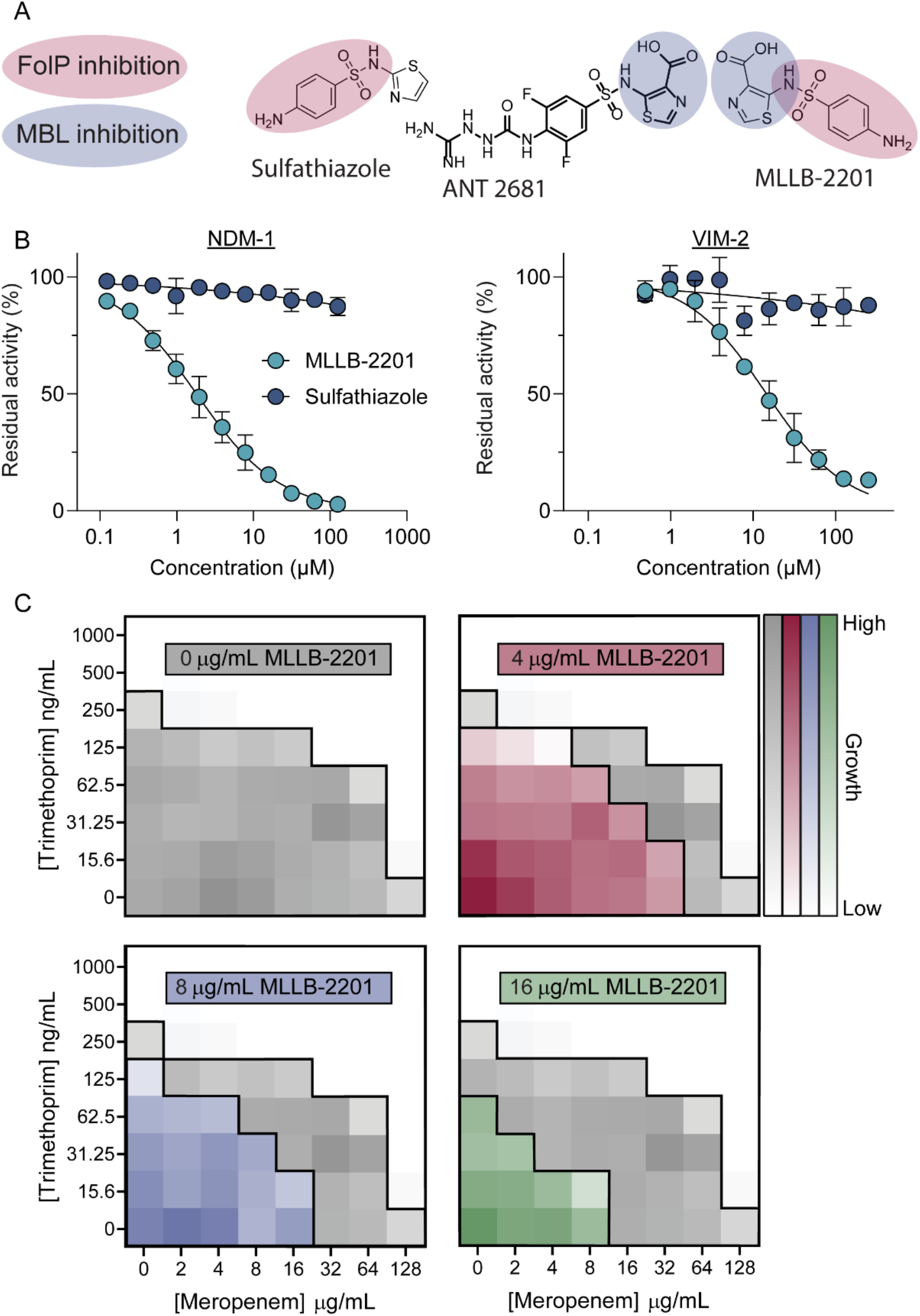
A dual-active inhibitor of metallo-β-lactamases overcomes resistance to meropenem via triple synergy. **A)** Chemical warheads responsible for FolP and metallo-β-lactamase inhibition are highlighted in red and blue, respectively. **B)** Effects of MLLB-2201 (light blue circles) and sulfathiazole (dark blue circles) treatment on NDM-1 (left) and VIM-2 (right) hydrolysis of meropenem. Increasing concentrations of each inhibitor are plotted on the X-axis, while the residual enzyme activity is plotted on the Y-axis as a percent of the uninhibited enzyme activity. Two biological replicates were performed, each with two technical replicates. A representative biological replicate is shown, with the mean of the technical replicates plotted as a coloured circle and the standard error of the mean plotted as error bars. To calculate IC50 values, curves were fitted to each data series and are shown as a black line. **C)** Three-way checkerboards showing the effects of trimethoprim, meropenem, and MLLB-2201 on growth of an *E. coli* strain expressing NDM-1. The concentrations of trimethoprim and meropenem are consistent across each checkerboard and the values are shown on the outer left and bottom axes, respectively. The concentration of the 3^rd^ compound, MLLB-2201, increases from left to right and top to bottom, and is indicated within each checkerboard. The growth values for the three triple-combination checkerboards are overlayed in coloured squares on top of the trimethoprim and meropenem-only checkerboard data to illustrate the effects of MLLB-2201 addition. The intensity of each colour indicates the relative growth where white represents no growth and full colour saturation represents full growth. Checkerboard assays were repeated in biological duplicates and data from a representative replicate are shown.

Leveraging this information, we synthesized MLLB-2201, a carboxylate-containing sulfonamide with potential dual activity against FolP and MBLs (**Fig 7a**). Gratifyingly, MLLB-2201 was a low micromolar inhibitor of the MBLs NDM-1 and VIM-2, with IC_50_ values of 1.8 and 14.5 µM, respectively (**Fig 7b**). Despite having a lower IC_50_ than ANT-2681^58^, MLLB-2201 still restored meropenem activity against an *E. coli* strain expressing NDM-1, whereas sulfathiazole could not (**Fig S8a**). Although treatment with MLLB-2201 alone resulted in minimal growth inhibition, it synergized with TMP against the Gram-positive pathogen, methicillin-resistant *Staphylococcus aureus* (**Fig S8b**), suggesting some FolP inhibition. Finally, when MLLB-2201 was tested in combination with TMP and meropenem against the *E. coli* strain expressing NDM-1, we observed three-way synergy (**Fig 7c**). MLLB-2201 (16 µg/mL) in combination with TMP (31.25 ng/mL) and meropenem (4 µg/mL) inhibited growth of this strain, representing 16-fold and 64-fold reductions in the single-antibiotic MICs.

## Discussion

Tetrahydrofolate plays a central role in one-carbon metabolism and the synthesis of critical metabolites, including thymidylate, purines, methionine, and glycine/serine interconversion. Given the importance of these metabolites for growth, disrupting folate biosynthesis is an effective strategy to kill pathogenic bacteria. The widespread dependence on folate metabolism for growth resulted in many interesting reports of potential connections between folate metabolism and broader cell physiology; however, the many consequences of folate inhibition that drive these observations also makes them challenging to study. Using bioinformatics, microscopy, and chemical genetics, we characterized the relationship between folate and PG metabolism in the important pathogen, *P. aeruginosa*. Our data point to a requirement for folate metabolism in maintaining cell envelope integrity as well as for full induction of *ampC* expression through the AmpR pathway.

Early hypotheses about connections between folate and PG metabolism date back about 50 years^11^, and likely stemmed from observation of morphological changes that followed SulA-mediated inhibition of cell division by induction of the SOS response. Investigation of the mechanism of mutual (rather than unidirectional) potentiation between TMP and SUL led to a model where these antibiotics were proposed to increase one another’s entry into the cell^60^.

Their synergistic mechanism is now known to arise from early folate pathway inhibition by a substrate accumulation feedback loop^61^, however data from the group that originally proposed increased compound uptake showed a striking effect of folate inhibition on the abundance of lipid II in *E. cloacae*^15^. Thus, while the initial model of mutual of potentiation was incorrect, the data supporting that model still revealed relevant effects of folate inhibition on the cell wall, which we further characterized here. This work also provides potential insight into the synergistic mechanisms of an antifolate-fosfomycin combination that was recently patented^22^.

Our data suggest that folate inhibition affects PG recycling. Using inactive point mutants of PBP4 and PBP5, we showed that overproduction of the sentinel anhMurNAc-pentapeptide recycling product that triggers AmpR-mediated expression of *ampC* led to TMP potentiation of non-inducing β-lactams. This potentiation also required AmpG, the muropeptide permease, and AmpR, suggesting that the sentinel product must enter the cytoplasm and activate the pathway by the canonical mechanism. Disrupting PG recycling in *P. aeruginosa* is also an effective strategy to potentiate FOS, as the recycling pathway in *P. aeruginosa* bypasses MurA^62,63^.

Antifolate-mediated disruption of PG recycling explains why TMP uniquely potentiated FOX and FOS in our initial PG-inhibitor interaction screen. Accordingly, we found that TMP significantly increased the abundance of GlcNAc-anhMurNAc and decreased the abundance of anhMurNAc. This finding supports the idea that TMP treatment disrupts PG recycling. Accumulation of stemless PG glycans is indicative of amidase activity, and the increase in GlcNAc-anhMurNAc coupled with a decrease in anhMurNAc suggests a possible bottleneck at the NagZ recycling step, either from reduced NagZ activity or increased production of stemless disaccharides. Alternatively, the lytic transglycosylase RlpA specifically cleaves stemless PG and liberates it for recycling^64^. Therefore, an increase in RlpA activity could also generate an excess of GlcNAc-anhMurNAc.

Despite defining the interaction of antifolates with specific PG inhibitors, we have yet to precisely identify which effects occurring downstream of folate inhibition are responsible for compromising PG metabolism. Our screen of the PA14 transposon library pointed to impacts on purine biosynthesis. The screen also identified mutants that were hypersensitive to TMP or FOS. Examination of purine mutants confirmed that *purC* and *purM* are hypersensitive to TMP, suggesting that small molecule inhibitors of these steps in purine synthesis could sensitize *P. aeruginosa* to the widely used TMP-SUL combination, cotrimoxazole. Previous work in *B. subtilis* showed that depleting purine but not thymidylate production (both folate dependent metabolites) caused a large decrease in PG turnover^65,66^. This observation aligns with the results of our mutant screen and the finding that DNA-synthesis inhibition is insufficient for FOS or FOX potentiation. It is unclear how the cells might sense purine depletion and concomitantly decrease the rate of PG turnover. Changes in the purine-dependent signalling nucleotide c-di-AMP affect PG synthesis in some bacteria^67^, but this second messenger has not yet been reported in *P. aeruginosa*. To uncover more factors involved in regulating PG turnover, the *dacBC*\*\* mutant that has constitutively elevated *ampC* expression could be used. Mutations that decrease *ampC* promoter activity in that background are likely to be compromised in PG recycling. Such a screen could uncover new factors affecting PG turnover, and point to novel ways to increase sensitivity of *P. aeruginosa* to AmpC-inducing β-lactams. Here we identified TMP as one such inhibitor that indirectly reduces degradation of β-lactams by AmpC, opening a new avenue for possible β-lactam adjuvant development.

Building upon the findings that folate inhibitors can potentiate β-lactams in *P. aeruginosa*, we designed MLLB-2201 as a dual inhibitor of metallo-β-lactamases and FolP, the target of sulfonamides. In theory, this strategy could achieve three-way potentiation with a β-lactam and TMP against *P. aeruginosa*. 1) TMP and MLLB-2201 synergize by inhibiting different steps in folate metabolism, similar to TMP-SUL. 2) MLLB-2201 blocks β-lactam degradation by metallo-β-lactamases. 3) TMP and MLLB-2201 decrease *ampC* induction, thus reducing β-lactam hydrolysis by AmpC. While we achieved some of these effects, *P. aeruginosa* remained largely unaffected by MLLB-2201 alone. A medicinal chemistry effort is currently underway to increase the potency of MLLB-2201, with a particular focus on improving FolP inhibition and anti-*Pseudomonas* activity. Together, this work demonstrates the potential for new, dual-target metallo-β-lactamase inhibitors with antibiotic and adjuvant properties.

Folate metabolism is vital to multiple aspects of bacterial physiology and there is an increasing recognition of the extent to which PG metabolism is coordinated with other cellular processes. For example, tailoring of PG peptide stems by D,D-carboxypeptidases, the targets of FOX, regulates activity of the Bam β-barrel assembly complex^68^. MurA and LpxC catalyze the first committed steps of PG and lipopolysaccharide (LPS) biosynthesis, respectively, and in *P. aeruginosa*, the FOS target MurA activates LpxC through a physical interaction that ensures the balanced consumption of UDP-GlcNAc by each pathway^69^. To coordinate positioning of the daughter chromosomes during cell division, SlmA interacts with and prevents polymerization of FtsZ^70^. Our data suggest that the folate and PG pathways are more intimately connected than previously appreciated, and that targeting their intersection could open new avenues for drug development.

## Materials and Methods

### Bacterial strains and growth conditions

All strains (**Table S1**) were stored at −80°C in 15% glycerol stocks that were used to inoculate overnight cultures. Overnight cultures were grown in 3 mL of lysogeny broth (LB, Lennox, Bioshop) at 37°C while shaking at 200 RPM. A 1:100 dilution of overnight cultures into fresh LB media was performed for subcultures. Antibiotics for plasmid maintenance were added when appropriate at the following concentrations: gentamicin at 15 µg/mL (Gent15) for *E. coli* or 30 µg/mL (Gent30) for *P. aeruginosa*; ampicillin at 100 µg/mL (Amp100) for *E. coli*; carbenicillin at 200 µg/mL (Carb200) for *P. aeruginosa*; 50 µg/mL kanamycin (Kan50) for *E. coli*.

### Plasmid and strain construction

pMS403 was constructed from pMS402Gm. pMS402Gm was created by digesting pMS402 and pPS856 with PstI, then isolating the gentamicin resistance cassette from the pPS856 digest reaction and using T4 ligase to insert the gentamicin cassette into pMS402. Next, pMS402Gm was digested at a BsiWI site within the *DHFRII* gene, the overhangs were filled in using the Klenow fragment polymerase, and the resulting blunt ends were rejoined by ligation with T4 ligase and transformed into *E. coli* DH5α. Transformants were selected on Gent15 and tested for trimethoprim sensitivity to determine *DHFRII* inactivation, then confirmed by sequencing.

pMS403(P*ampC*) was created by PCR amplifying the *ampC* promoter with P*ampC* Fwd and P*ampC* Rvs (all primers are listed in **Table S1**), then digesting both pMS403 and the purified PCR product with BamHI and XbaI, and ligating the two purified digest products with T4 ligase. The ligation product was transformed into *E. coli* DH5α and plasmids isolated from Gent15 resistant colonies were verified for the correct insert with sequencing.

pUCP20 (*DHFRII*) was created by PCR amplifying the *DHFRII* gene from the pMS402 backbone with *DHFRII* Fwd and *DHFRII* Rvs primers. Next, pUCP20 and the purified PCR product were digested with EcoRI and HindIII and the purified digest products were ligated together with T4 ligase, then transformed into *E. coli* DH5α. Plasmids from Amp100 resistant colonies were isolated and sequenced to verify the correct insert.

pEX18Gm (*ampC*), pEX18Gm (*ampC* S90A), and pEX18Gm (*ampG*) were constructed by digesting the gBlock inserts in pUC57 (Genscript) with EcoRI and HindIII (or SacI for *ampG*). The inserts were gel purified and ligated using T4 ligase into empty pEX18Gm that was digested with EcoRI and HindIII/SacI. The ligations were transformed into *E. coli* DH5α and transformants were selected on Gent15 plates. Plasmids isolated from the transformants were sequenced to verify the correct insertion.

pEX18Gm (*ampR*) was constructed by amplifying 500 bp regions of chromosomal DNA from PAO1 that flank the 5’ and 3’ ends of *ampR* using two PCR reactions containing the d*ampR* Up Fwd and d*ampR* Up Rvs primers, or the d*ampR* Dwn Fwd and d*ampR* Dwn Rvs primers. The purified PCR products from these reactions were combined and used as templates for overlap extension PCR with the d*ampR* Up Fwd and d*ampR* Dwn Rvs primers, and the purified PCR product was digested with BamHI and HindIII. The purified digest product was ligated into linearized pEX18Gm (digested with BamHI and HindIII) using T4 ligase. The ligations were transformed into *E. coli* DH5α and transformants were selected on Gent15 plates. Plasmids isolated from the transformants were sequenced to verify the correct insertion.

pMS403 and pUCP20-based plasmids were introduced to *P. aeruginosa* by electroporation. pEX18Gm-based plasmids were first introduced into *E. coli* SM10 by electroporation, then *E. coli* SM10 was used to transfer the plasmid into *P. aeruginosa* by conjugation. *P. aeruginosa* cells containing the plasmid were selected for on *Pseudomonas* isolation agar (PIA) containing 100 µg/mL of gentamicin to select for the first recombination event. Colonies from the PIA Gent100 plates were streaked onto LB 5% sucrose no NaCl agar plates and incubated at 30°C to select for a second recombination event. Colonies from the LB sucrose plates were patched onto LB and LB Gent30 plates. Patches that grew on LB but not LB Gent30 plates were tested for loss of intrinsic ampicillin resistance, which indicates loss of AmpC activity. Mutants were confirmed with PCR.

Strains containing the *dacB*/dacC\** catalytically inactive point mutations were constructed by site-directed mutagenesis of PAO1 wild-type alleles. The *dacB*\* primer included a TCG→GCG mutation that converted the serine 72 codon to an alanine, while the *dacC*\* primer included an AGC→GCG mutation that converted the serine 64 codon to an alanine. These mutant alleles were crossed into the chromosome of PAO1 using allelic exchange with the pEX18Gm plasmid. Mutants were confirmed by sequencing mutant alleles amplified with PCR.

### Determination of minimal inhibitory concentration

Minimum inhibitory concentration (MIC) assays were performed by passaging overnight cultures into 3 mL of 10:90 LB (1:9 ratio of LB to phosphate buffered saline) and growing to an OD_600_ of ∼0.1-0.3. Cultures were normalized to an OD_600_ of 0.1 and diluted 1:500 into 10:90 LB. Two microliters of the indicated antibiotic were added to the wells in rows A-F of a 96-well, and 2 µL of the vehicle control was added to the wells in rows G and H. Antibiotics were added at 75x the final concentration. After the antibiotics were added, 148 µL of the diluted, normalized culture was added to the wells in rows A-G. Row H served as a sterility control and 148 µL of sterile 10:90 was added to these wells. Plates were incubated at 37°C while shaking for 18 hours. Growth (OD_600_) was measured using a plate reader (Multiskan Go, Thermo Fisher Scientific).

### Checkerboard assays

Overnight cultures were diluted 1:100 into 3 mL of LB and grown to ∼0.1-0.3 OD_600_. For PA14 transposon mutants, overnight cultures were made using LB Gent30. Subcultures were normalized to 0.1 OD_600_ and diluted 1:500 in fresh LB. Serial dilutions of 1 µL of one antibiotic was added in decreasing concentrations across rows A-H for columns 3-10, while the second antibiotic was added in decreasing concentrations across columns 3-10 for rows A-H. Two microliters of the respective vehicle control were added to columns 1, 2, 11, and 12. The 1:500 diluted cultures were added to the wells to a final volume of 150 µL. Sterility wells were filled to 150 µL with sterile LB. Plates were incubated at 37°C for 18 h while shaking at 200 rpm. Growth (OD_600_) was measured using a plate reader (Multiskan Go, Thermo Fisher Scientific).

### Bioinformatic analysis

We established a local blast database by compiling a comprehensive genome dataset from the GTDB, encompassing a total of 47,894 genomes as of June 2023. Subsequently, we conducted tblastn searches on this local GTDB database^71^, comparing two protein sequences, *folP* and *glmM*, from PAO1. To identify the best matches, we analyzed the tabular blast output in R v4.2.0, selecting top hits based on their percent identity. We also removed duplicate hits within contigs and calculated the genomic distance between the starting point of *glmM* and the ending point of *folP*.

To test whether there is a potential phylogenetic association between *folP* and *glmM*, we focused on a specific subset of proteobacterial genomes from our database. Specifically, we targeted proteobacterial families with a representation of 100 or more genomes labeled as complete in the RefSeq database. This subset, encompassing a total of 939 genomes, was then used to construct a phylogenetic tree. This tree was generated through sequence alignment of GTDB markers and subsequent approximately maximum likelihood tree construction using FastTree 2^72^. The results were visually represented using the ggplot package in R v4.2.0, and the phylogenetic tree was further enhanced and visualized using iTOL^73^.

### Luminescent *ampC* promoter-reporter assay

Overnight cultures of PAO1 containing pMS402(Empty) or pMS402(P*ampC*) were made by inoculating 3 mL of LB Gent30 from frozen stocks and were incubated at 37 °C with shaking. Subcultures were made by transferring 120 µL of the overnights into 3 mL of LB Gent30 and incubated at 37°C with shaking for ∼2 hours until an OD_600_ of ∼0.1-0.3 was reached. Cultures were normalized to an OD_600_ of 0.1, then diluted 1:500 in fresh LB Gent30. Assays were prepared in white-walled 96-well plates with clear bottoms (Corning). Two microlitres of dilutions of trimethoprim were added across rows A-E at the indicated concentrations. Cefoxitin was added to the wells in rows A-F at a final concentration of 50 µg/mL. A DMSO vehicle control was added to wells in rows G and H. Afterwards, 148 µL of the diluted culture was added to each well of the plate, except row H, which served as a sterility control and received 148 µL of LB Gent 30. The plate was incubated at 37°C with continuous double orbital shaking for 16 hours in a Synergy Neo (Biotek) plate reader. Growth (OD_600_) and luminescence (luminescence fiber) measurements were taken every 15 minutes and promoter activity at 8 hours was graphed. Relative luminescence units (RLU) were calculated by dividing each well’s luminescence value by its growth at the corresponding time point.

### Determination of β-lactamase activity

Whole cell AmpC activity was determined as previously described with minor modifications^74^. Briefly, overnight cultures were grown in LB and subcultures were made by diluting the overnight culture 1:25 into fresh LB, then incubated for 2 hours at 37°C while shaking. Cells were then passaged again into fresh media containing 50 µg/mL of cefoxitin or a DMSO vehicle control, and trimethoprim at the indicated concentrations. The cultures were then incubated again for ∼2-3 hours until cells reached OD6_600_ 0.4-0.6, then the cultures were normalized to 0.3 OD_600_ and 1 mL of normalized culture was centrifuged at 21 000 x g for 1 minute. The supernatant was decanted, and the cell pellet was washed with 1 mL of 50 mM sodium phosphate buffer (pH 7.4), then resuspended in 1 mL of the sodium phosphate buffer.

Cells placed on ice then lysed with sonication by a microtip (Sonicator 2000, Microsonix) using three 10 second pulses with 10 second pauses between pulses. The cell debris was pelleted by centrifugation at 21 000 x g for 5 minutes, and 500 µL of supernatant was moved to a fresh Eppendorf tube. Five microliters of nitrocefin were added to wells of a clear flat-bottom 96 well plate for a final concentration of 50 µM, then 195 µL of the clarified cell lysate was added to each well. Absorbance at 490 nm was read with a spectrophotometer (Multiskan Go, Thermo Fisher) immediately and for every 15 seconds thereafter until the linear range of the uninhibited positive control was exceeded.

*In vitro* AmpC β-lactamase activity was measured with purified commercial AmpC (Sigma Aldrich) from *P. aeruginosa*. AmpC was diluted to 400 nM (final concentration) across a serial dilution of the indicated compound in 50 mM sodium phosphate buffer (pH 7.4), then added to the wells of a 96 well plate containing nitrocefin at 50 µM (final concentration) to a final volume of 200 µL. Then, absorbance at 490 nm was read immediately and for every 15 seconds after for 10 minutes by a spectrophotometer (Multiskan Go, Thermo Fisher). The data within the time points for the linear range of the no compound AmpC control were used for analysis.

*In vitro* metallo β-lactamase activity was measure using NDM-1 (5 nM) or VIM-2 (50 nM) incubated in reaction buffer (25 mM HEPES-NaOH, 10 μM ZnSO_4_, pH 7.5) containing varying amounts of inhibitor (500 – 1 μM) and incubated for 5 min at room temperature. Residual enzyme activity was determined by measuring β-lactam hydrolysis spectrophotometrically at 300 nm by adding a saturating amount of meropenem (500 μM) to the reaction mixtures containing enzyme and inhibitor. β-Lactamase assays were performed in a clear flat-bottom 96-well plate at 25°C with a final assay volume of 200 μL and monitored with a BioTek Synergy H1 microplate reader over 10 min. All reactions were performed in duplicate unless otherwise stated.

### PA14 library screen

Rectangular plates with 25 mL of LB 1.5% agar were poured that contained no antibiotic, 32 µg/mL fosfomycin, 32 µg/mL trimethoprim, or both antibiotics, and allowed to dry overnight. An ordered PA14 transposon mutant library was pinned from source plates in 1536 colony density onto the rectangular agar plates using a ROTOR HDA robotic colony replicator (Singer Instruments). The screen was performed in duplicates using different source plates. Plates were incubated for 18 h at 37°C, then imaged using a Phenobooth imaging system (Singer Instruments) using the transmissive light mode. Images were further processed in FIJI (Image J) as described previously^75^. Briefly, the light absorbed by each colony was converted into an integrated density value. Integrated densities were then normalized for plate position effects and the relative growth was determined by comparing to the untreated control.

### Soluble muropeptide extraction and analysis

Soluble muropeptides were prepared according to Weaver *et al.* (2022) with minor modifications^76^. Briefly, 50 mL flasks of LB were inoculated with 0.5 mL of an overnight culture. Flasks were prepared in technical duplicate with antibiotics added at the indicated concentrations. Cultures were incubated at 37°C while shaking until an OD_600_ of ∼0.4 was reached. Then, flasks were immediately placed on ice, cultures were normalized to an OD_600_ of 0.3, and 40 mL were transferred to pre-chilled 50 mL falcon tubes. The normalized cultures were then centrifuged at 5030 x g and 4°C for 20 minutes on an Avanti J-26 XPI (Beckman-Coulter) centrifuge (JS 5.3 rotor). The supernatant was decanted, and cell pellets were resuspended in 1 mL ice cold 0.9% NaCl, washed twice with 1 mL 0.9% NaCl, and finally resuspended in 1 mL of sterile nuclease free water. The Eppendorf tubes containing the resuspended cells were boiled for 30 minutes to lyse the cells and centrifuged for 15 minutes at 21 000 x g in a benchtop centrifuge to pellet the cell debris. The supernatant was then passed through a 0.2 µm filter and frozen at −80°C. Samples were concentrated as needed under vacuum using a lyophilizer (Virtis). Lyophilized samples were dissolved in sterile nuclease free water and the pH was adjusted to ∼3 using formic acid. After adjusting the pH, samples were centrifuged for 10 minutes at 21 000 x g in a benchtop centrifuge to pellet any precipitate.

Ten microliters of each sample were injected into an LC/Q-TOF (Agilent 6546) and separated using an Eclipse Plus C18 column (Agilent, 95 Å pore size, 2.1x×00 mm, 1.8 µm) at 50°C. Separating of PG species was achieved using a linear gradient of buffer A (water+0.1% formic acid) to buffer B (acetonitrile+0.1% formic acid) over a 56 minute run time with a 0.4 mL/minute flow rate. The Q-TOF instrument was run in negative ionization mode with the following parameters: 4000 V capillary voltage, 300°C source temperature, 300°C sheath gas temperature, and a scan range of 100-1700 *m/z*. Data acquisition and analysis was performed using the Agilent MassHunter qualitative analysis (v10.0) software. Extracted ion chromatograms were manually curated using theoretical muropeptide masses. The selected muropeptides were based on *m/z* values of 595.6639 for UDP-MurNAc-AEmAA (doubly charged species), 524.6263 for UDP-MurNAc-AEm (doubly charged species), 477.1726 for GlcNAc-anhMurNAc (singly charged species), and 274.0932 for anhMurNAc (singly charged species).

### Western blots

Quantification of total AmpC protein levels by western blot was performed exactly as described in Lamers et al.^77^. Samples were prepared identically, with the strains and conditions described in the figure caption. The non-specific band shown in the image of the blot was used as a loading control.

### Flux balance analysis

The iPAO1 genome scale metabolic model developed by Zhu *et al*.^49^ was imported into Matlab R2020a (MathWorks) and the Cobra toolbox (Version 3.0)^48^ was used to perform FBA with a Gurobi mathematical solver (Version 9.0.2). FBA-Div was used to simulate antibiotic treatment^78^, where the substrates of inhibited reactions are diverted to a waste reaction. The iPAO1 model contains an irreversible and reversible reaction for FolP; therefore, to inhibit both reactions, the irreversible reaction (rxn02201) was removed from the model and flux through the reversible reaction (rxn02200) was reduced, and the substrate (dihydropteroate) was diverted to a waste reaction. To simulate a dilution range of antibiotic treatment, the optimal flux for each reaction was determined under no reaction inhibition, then the flux rate was simulated at 20% intervals from 0-100% of the optimal flux rate. The calculated biomass production rate at the steady state for each interval was determined in single and double reaction inhibitions to generate growth values that were used to create a checkerboard. Rich media growth conditions with an abundance of nutrients was assumed for the metabolic modelling.

### Microscopy

Overnight cultures were diluted 1:100 in LB with or without the trimethoprim at the indicated concentrations and incubated until an OD_600_ of ∼0.4. All cultures were normalized to an OD_600_ of 0.3, then 1 mL of each culture was pelleted, washed with 1 mL of sterile PBS, then resuspended in 100 µL of sterile PBS. Four microliters of the resuspended cells were spotted on 1.5% M9+glucose agarose pads on glass slides and covered with coverslips. Cells were imaged with a Nikon Ti-2 Eclipse inverted confocal microscope using a 60X oil immersion objective lens. Identical settings were used for all micrographs captured.

### Synthesis of MLLB-2201

See supplemental information.

### Construction of graphs, structures, and statistical analysis

All graphs and checkerboards were created using GraphPad (Prism, Version 10), and statistical analyses were performed using GraphPad as well. The graph in Figure 1C was created using R and the tree in Figure 1D was created using iTOL. The structures shown in Figure 1A were downloaded from the protein data bank and modelled in ChimeraX (Version 1.4)^79^.

## Supporting information

Yaeger et al. Extended Data Figs 1-7

Yaeger et al. Supplementary Data Synthesis of MLLB-2201

Yaeger et al. Supplementary Table S1

Yaeger et al. Extended Data Movie 1

## Acknowledgements

This work was funded by Natural Sciences and Engineering Research Council of Canada (www.nserc-crsng.gc.ca) grant RGPIN-2021-04237 to LLB and by infrastructure funding from Canada Foundation for Innovation and the Ontario Research Fund (ORF-RE09-047). LLB holds a Tier I Canada Research Chair in Microbe-Surface Interactions (CRC-2021-00103). LNY holds an NSERC PGS-D award. OG holds a Chercheur Boursier Junior 1 Fellowship, from the Fonds de Recherche du Québec-Santé.

## Notes

### Competing Interest Statement

The authors have declared no competing interest.

