## Supplementary material for "Metabolic connections between folate and peptidoglycan pathways in *Pseudomonas aeruginosa* inform rational design of a dual-action inhibitor": Yaeger et al. Extended Data Figs 1-7

TMP Treated Time Course.avi

**Extended Data Movie 1. TMP treatment causes explosive cell lysis.** A time lapse of TMP treated cells was created by capturing images every 15 seconds, compiling all images into an image stack, and exporting the stack as an .avi file at 5 frames per second. The round cell undergoes explosive cell lysis.

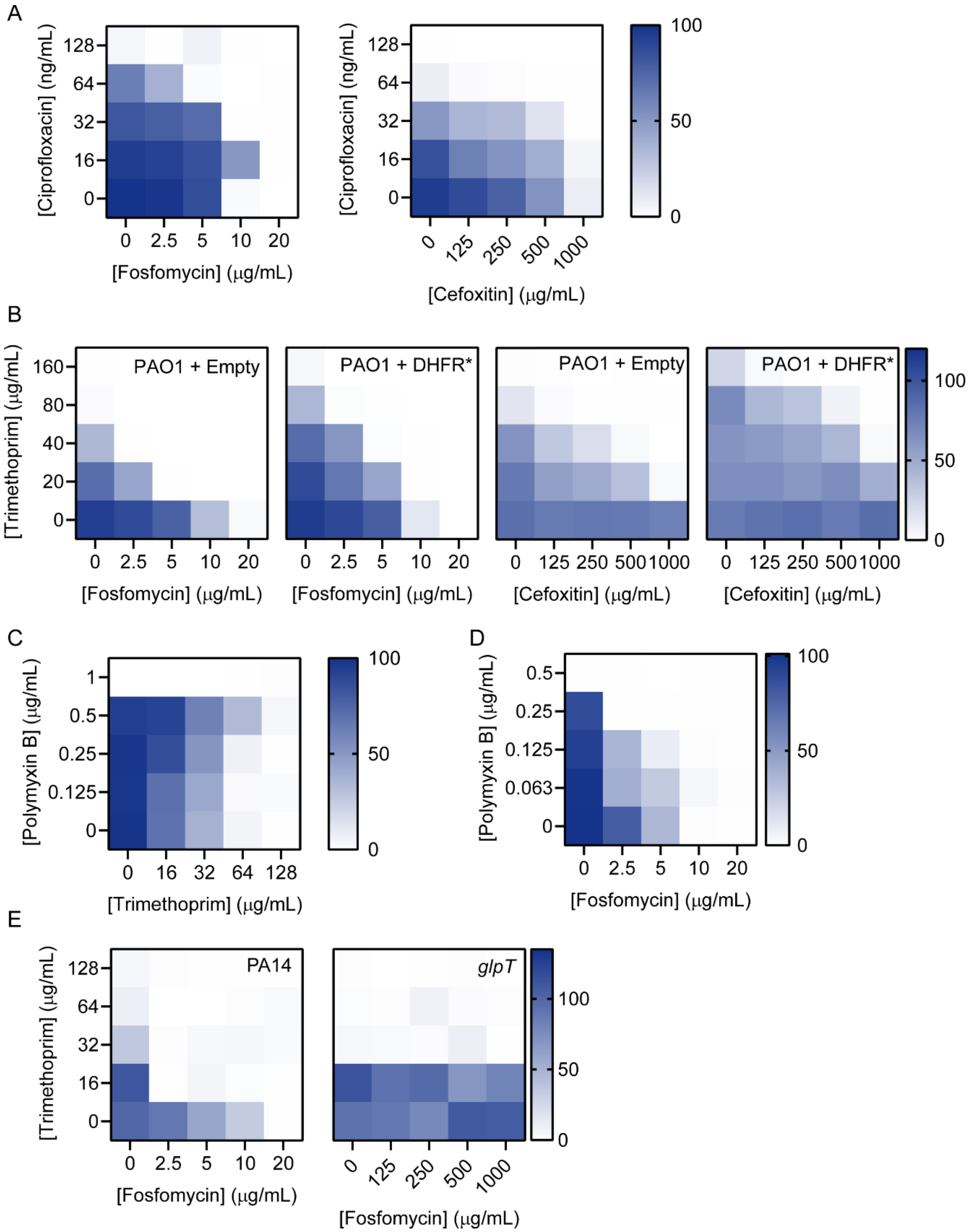
A
[Ciprofloxacin] (ng/mL)
128
64
32
16
0
0
2.5
5
10
20
[Fosfomycin] (μg/mL)
[Ciprofloxacin] (ng/mL)
128
64
32
16
0
0
125
250
500
1000
[Cefoxitin] (μg/mL)
100
50
0
B
[Trimethoprim] (μg/mL)
160
80
40
20
0
PAO1 + Empty
PAO1 + DHFR*
PAO1 + Empty
PAO1 + DHFR*
0
2.5
5
10
20
[Fosfomycin] (μg/mL)
0
2.5
5
10
20
[Fosfomycin] (μg/mL)
0
125
250
500
1000
[Cefoxitin] (μg/mL)
0
125
250
500
1000
[Cefoxitin] (μg/mL)
100
50
0
C
[Polymyxin B] (μg/mL)
1
0.5
0.25
0.125
0
0
16
32
64
128
[Trimethoprim] (μg/mL)
100
50
0
D
[Polymyxin B] (μg/mL)
0.5
0.25
0.125
0.063
0
0
2.5
5
10
20
[Fosfomycin] (μg/mL)
100
50
0
E
[Trimethoprim] (μg/mL)
128
64
32
16
0
PA14
glpT
0
2.5
5
10
20
[Fosfomycin] (μg/mL)
0
125
250
500
1000
[Fosfomycin] (μg/mL)
100
50
0

**Extended Data Figure 1. TMP potentiates FOS and FOX through its primary MOA. A-E)** Checkerboard assays between two antibiotics (labelled on X- and Y-axes). Checkerboards were performed using an 8x8 concentration range and condensed into a 5x5 concentration range (labelled on axes) for each figure. The scale bar on the right indicates the amount of growth where white shows no growth and dark blue shows the highest growth. Growth is represented as a percent of the untreated control growth for each matched assay. Each assay was performed in biological duplicate and a representative replicate is shown. For **B and E)**, the strains used for each assay are labelled in the top right corner. PAO1 was used for **A, C, and D**.

2

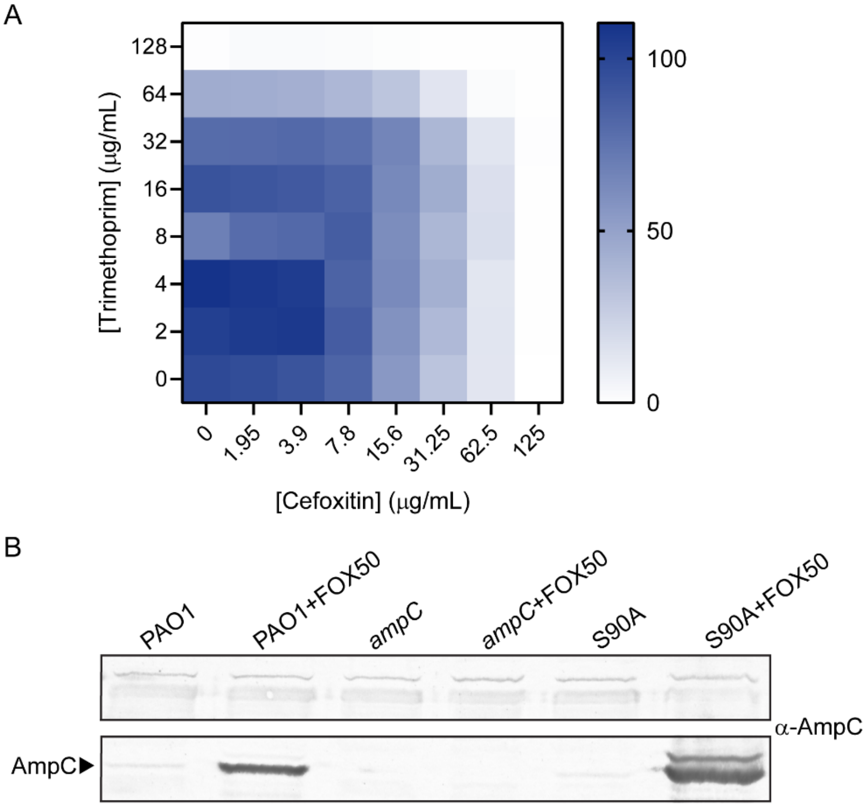


**Extended Data Figure 2. Induction of AmpC is insufficient for TMP potentiation of FOX. A)** An 8x8 checkerboard assay measuring interactions between trimethoprim and cefoxitin using the *ampC* S90A strain. The colour gradient from white to blue indicates the growth as a percent of the untreated control and the legend is shown on the right. Assays were performed in biological duplicate and a representative replicate is shown. **B)** A western blot for AmpC from whole cell lysates. Samples were prepared from cultures of the indicated strain and growth condition above each lane. FOX50 indicates 50 μg/mL of cefoxitin. The bottom box shows the bands corresponding to the molecular mass of AmpC (indicated with the arrowhead on the left). Note that the second, slightly larger band for the S90A+FOX50 lane is likely unprocessed cytoplasmic AmpC that still contains the N-terminal signal peptide due to high levels of induction. The top box shows non-specific bands that bind the AmpC antibody and are used as a loading control. The western blot was repeated using two independently prepared samples as biological replicates, and a representative blot is shown.

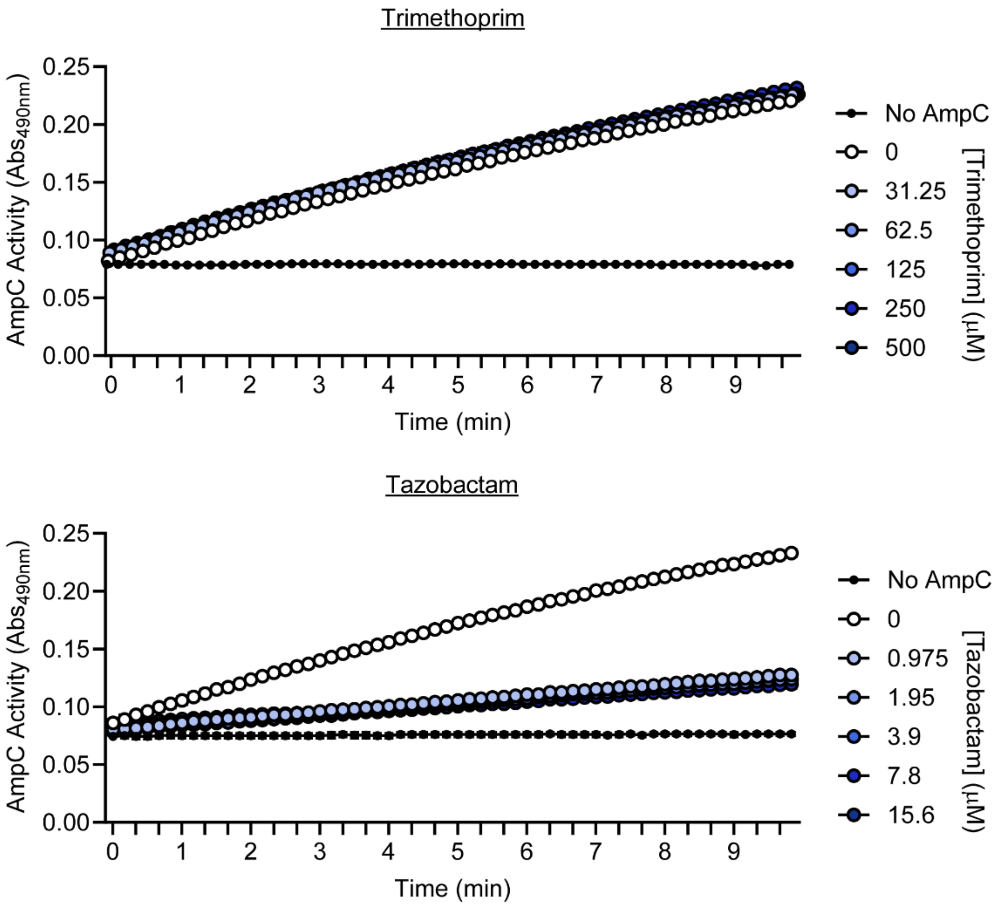


**Extended Data Figure 3. Trimethoprim does not directly inhibit AmpC.** The effect of trimethoprim (top) and the β-lactamase inhibitor Tazobactam (bottom) on purified AmpC activity was measured across time (X-axis) using nitrocefin hydrolysis. Hydrolyzed nitrocefin absorbs light at 490 nm and absorbance values are plotted along the Y-axis. Concentrations of each compound are shown on the legend on the right. Circles indicate the mean of two technical replicates, and the error bars indicate the standard error of the mean. Experiments were repeated in biological duplicate and a representative replicate is shown.

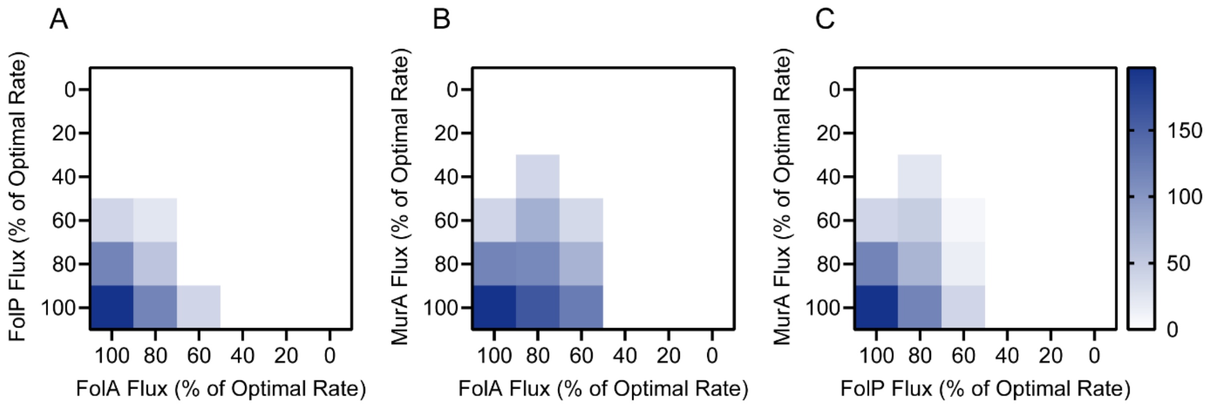


**Extended Data Figure 4. Antifolate-fosfomycin interactions are not predicted by *in silico* modelling. A-C)** Heatmaps showing the combinatorial effects of reducing flux through FolP, FolA, and MurA in an *in silico* genome scale metabolic model. The axes show the flux through each enzymes reaction for each simulation as a percent of the optimal steady state flux rate. The colour indicates the final biomass production rate at each flux rate coordinate on the plot, where white indicates no biomass production rate, and dark blue indicates the highest biomass production rate (legend shown on the right).

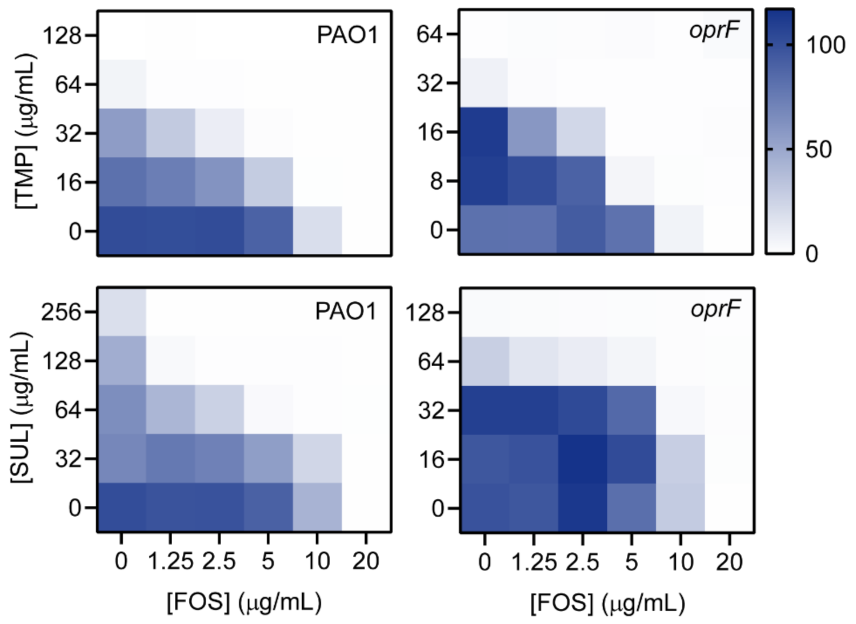


**Extended Data Figure 5. Loss of OprF is redundant with FOS potentiation.** Checkerboard assays were performed with an 8x8 grid and condensed to a 5x6 grid. The antibiotics used in each assay are shown on the axes, where the top two heatmaps used TMP and FOS, while the bottom two used SUL and FOS. The strain used for each assay is indicated in the top right corner. Note that the *oprF* mutant is more sensitive to antifolates, so a lower concentration range is shown. Growth was calculated as a percent of the untreated control and was plotted as a colour gradient where white indicates no growth and dark blue indicates the highest growth (legend shown in the top right). The FICIs for TMP-FOS are 0.5 and 0.625 for PAO1 and *oprF*, and for SUL-FOS are 0.375 and 0.625 for PAO1 and *oprF*. Checkerboards were repeated in biological duplicate and a representative replicate is shown.

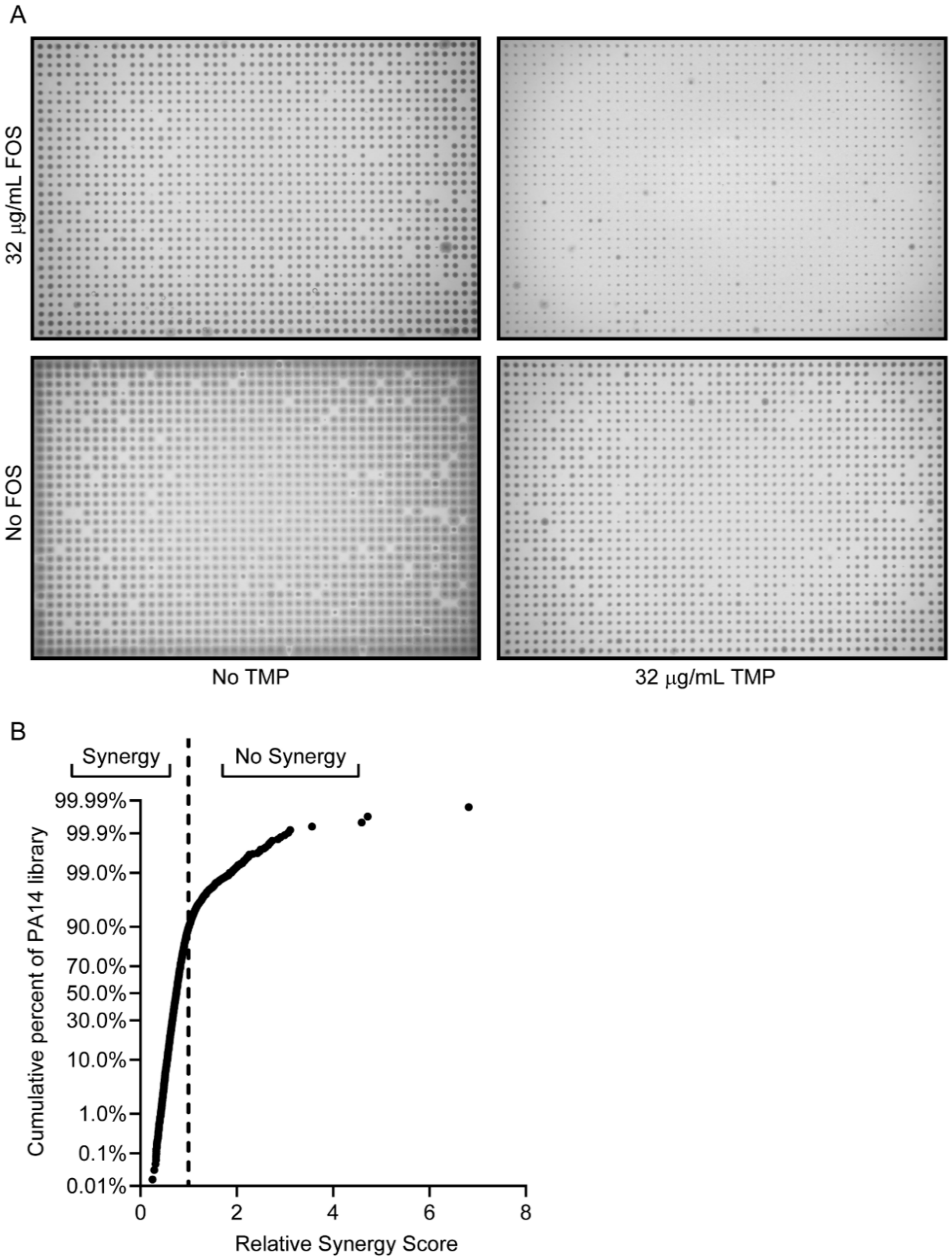


**Extended Data Figure 6. A 1536-colony density chemical genetic screen identifies antibiotic interaction determinants. A)** Scans of agar plates with 1536 colonies, where each colony is a PA14 transposon mutant. The conditions of each plate correspond to the labels on the left and bottom, where the top two plates have FOS supplemented and the right two plates have TMP supplemented. **B)** A dot plot showing the distribution of synergy across the PA14 transposon library. The cumulative percentage of the library at or below a given synergy score is plotted on the Y-axis, while the synergy score is plotted on the X-axis. The dashed line indicates the cutoff for synergy. Synergy occurs in over 90% of the mutants. Note that single antibiotic hypersensitive mutants are likely to score as lacking synergy, which will overrepresent the frequency of mutants lacking synergy.

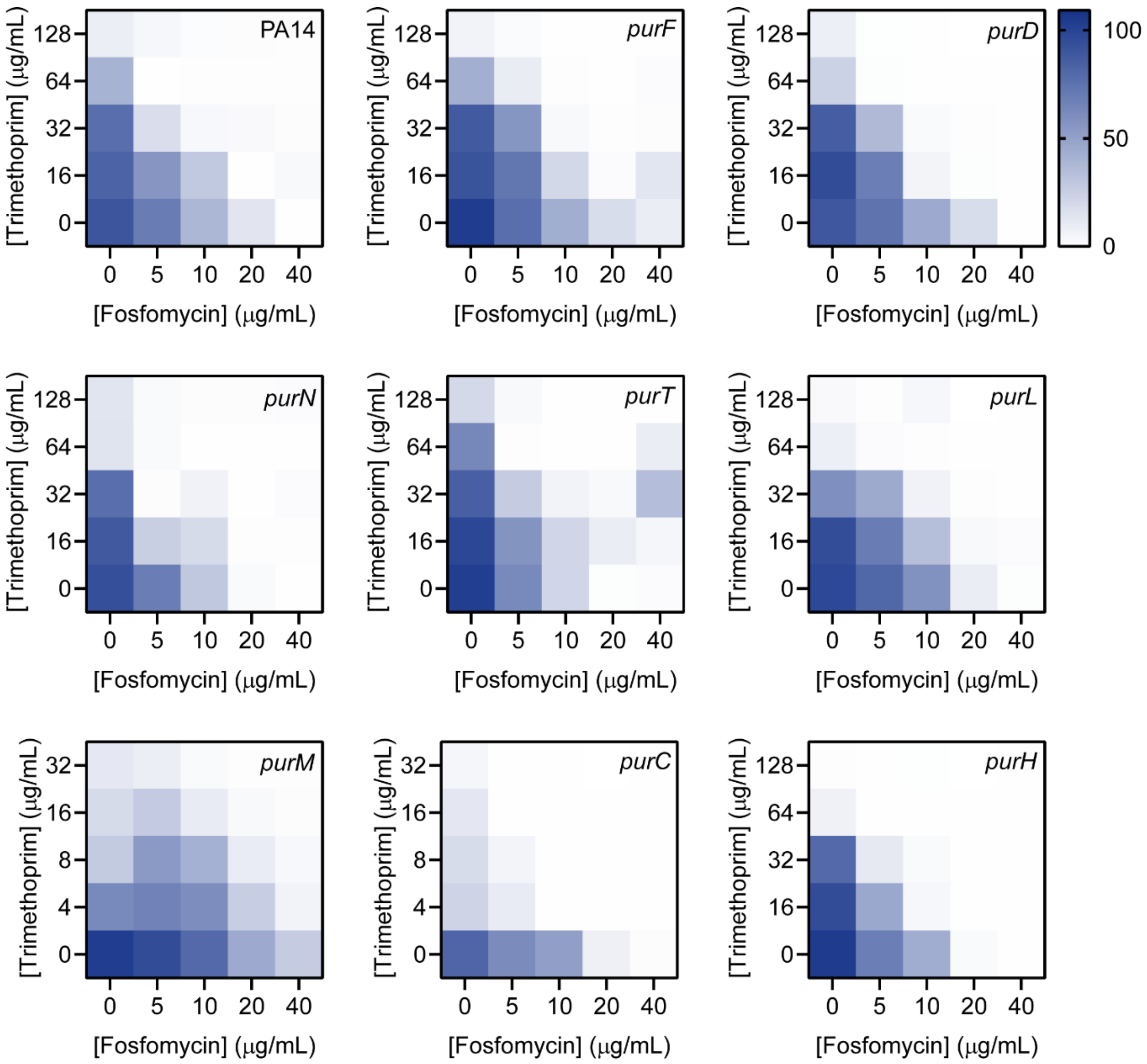


**Extended Data Figure 7. Effects of disrupting purine biosynthesis on the TMP FOS interaction.** Heatmaps showing 5x5 checkerboards that condense data from 8x8 checkerboards. The intensity of the blue colour corresponds to the growth as a percent of the vehicle control shown in the legend beside the top right heatmap. The strain used in each assay is indicated in the top right corner. The correct transposon insertion for each *pur* mutant was validated by colony PCR using the corresponding primers in Table S1. The FICIs are as follows: PA14 = 0.5; *purF* = 0.5; *purD* = 0.375; *purN* = 0.5; *purT* = 0.625; *purL* = 1; *purM* = <0.5; *purC* = 0.375; *purH* = 0.625. Each checkerboard was repeated in biological triplicate, and the mean growth at every matched antibiotic combination was plotted.
