## Supplementary material for "Metabolic connections between folate and peptidoglycan pathways in *Pseudomonas aeruginosa* inform rational design of a dual-action inhibitor": Yaeger et al. Supplementary Data Synthesis of MLLB-2201

**Experimental procedures for the synthesis of MLLB-2201:**

**General:**

Chemical shifts in $^{1}$H NMR and $^{13}$C NMR spectra are reported in parts per million (ppm) relative to tetramethylsilane (TMS), with calibration of the residual solvent peaks according to values reported by Gottlieb et al. (chloroform: $\delta_H$ 7.26, $\delta_C$ 77.16; DMSO: $\delta_H$ 2.50, $\delta_C$ 39.52).[1] When peak multiplicities are given, the following abbreviations are used: s, singlet; d, doublet; t, triplet; q, quartet; sept., septet; dd, doublet of doublets; m, multiplet; br, broad; app., apparent; *gem*, geminal. $^{1}$H NMR spectra were acquired at 400 or 700 MHz with a default digital resolution (Brüker parameter: FIDRES) of 0.22 and0.15 Hz/point, respectively. Coupling constants reported herein therefore have uncertainties of ±0.4 Hz and ±0.3 Hz, respectively. Synthetic experimental procedures were adapted from work previously published by Davies et al.[2] Reactions were carried out at room temperature (rt) if temperature is not specified. Compounds purified by normal-phase flash chromatography used Teledyne Combi*Flash* Rf+ and NextGen 300+ purification systems (www.teledyneisco.com) equipped with pre-packed silica cartridges (either 40–60 μM or 20–40 μM particle size). High-performance liquid chromatography was conducted using an Agilent 1290 Infinity II Preparative HPLC system and an eluent system comprising of 5mM ammonium acetate buffer and acetonitrile. Low-resolution mass spectral (LRMS) measurements were recorded on an Advion Expression CMS Compact Mass Spectrometer (Albany, NY). High-resolution mass spectrometric (HRMS) data was obtained using an Brüker micrOTOF II system with electrospray ionization (ESI) and paired with an Agilent HPLC and UV detector.

Reagents were purchased from Ambeed, Combi-Blocks, Fisher Scientific, and Sigma Aldrich, and used without any additional purification.

---

[1] Gottlieb, H. G.; Kotlyar, V.; Nudelman, A. NMR Chemical Shifts of Common Laboratory Solvents as Trace Impurities. *J. Org. Chem.* **1997**, *62*, 7512–7515.

[2] Davies, D.T.; Leiris, S.; Sprynski, N.; Castandet, J.; Lozano, C.; Bousquet, J.; Zalacain, M.; Vasa, S.; Dasari, P.K.; Pattipati, R.; Vempala, N.; Gujjewar, S.; Godi, S.; Jallala, R.; Sathyap, R.R.; Darshanoju, N.A.; Ravu, V.R.; Juventhala, R.R.; Pottabathini, N.; Sharma, S.; Pothukanuri, S.; Holden, K.; Warn, P.; Marcoccia, F.; Benvenuti, M.; Pozzi, C.; Mangani, S.; Docquier, J.D.; Lemonnier, M.; and Everett, M. ANT2681: SAR Studies Leading to the Identification of a Metallo-β-lactamase Inhibitor with Potential for Clinical Use in Combination with Meropenem for the Treatment of Infections Caused by NDM-Producing Enterobacteriaceae. *ACS Infectious Diseases*, **2020**, 6 (9), 2419-2430.

**Ethyl 5-((4-nitrophenyl)sulfonamido)thiazole-4-carboxylate**

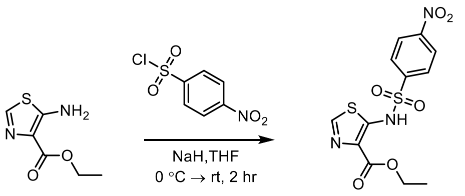


To a clean, dry 50 mL round bottom flask, sodium hydride (60% in oil, 66.9 mg, 1.74 mmol, 3 equiv.) was suspended in dry tetrahydrofuran (THF). The mixture was cooled to 0 °C before the addition of ethyl 5-amino-thiazole-4-carboxylate (100.0 mg, 0.58 mmol, 1 equiv.) and stirred for at least ten minutes. Afterward, 4-nitrobenzenesulfonyl chloride (154.4 mg, 0.69 mmol, 1.2 equiv.) was added at 0 °C and the mixture was left to warm to room temperature and stir for 2 hours. Reaction progress was monitored with thin layer chromatography (TLC) [5% methanol (MeOH): dichloromethane (DCM)]. The reaction was quenched with an aqueous solution of saturated ammonium chloride and diluted with diethyl ether. The resulting precipitate was then collected via vacuum filtration and washed with multiple aliquots of diethyl ether and water. The product was dried under vacuum to obtain a brownish-yellow solid. The crude product was purified using normal-phase chromatography on silica gel (0→20% MeOH/DCM) to afford the sulfonamide as an orange-yellow solid (114.0 mg, 0.31 mmol, 55 %). If desired, the crude product could be carried forward to the next reaction without any prior purification.

$R_f$ = 0.31 (5% MeOH/DCM).

$^1$H NMR (400 MHz, DMSO-$d_6$) δ 8.70 (s, 1H), 8.39 (d, J = 8.9 Hz, 2H), 8.04 (d, J = 8.9 Hz, 2H), 4.14 (q, J = 7.1 Hz, 2H), 1.17 (t, J = 7.1 Hz, 3H).

$^{13}$C NMR (100 MHz, DMSO-$d_6$) δ 159.47, 148.31, 146.27, 143.20, 131.15, 126.74, 122.98, 58.88, 12.38.

LRMS m/z: $[M + H]^+$ calculated for $C_{12}H_{12}N_3O_6S_2^+$ 358.0162 ; Found 358.1500.

HRMS (ESI) m/z: $[M - H]^-$ calculated for $C_{12}H_{10}N_3O_6S_2^-$ 356.0017; Found 356.0020.

**Ethyl 5-((4-aminophenyl)sulfonamido)thiazole-4-carboxylate**

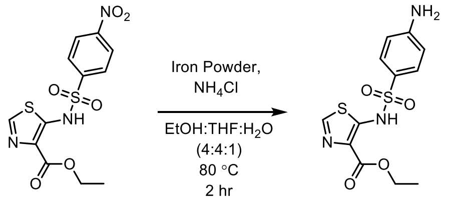


In a clean, dry 50 mL round bottom flask, Ethyl 5-((4-nitrophenyl)sulfonamido)thiazole-4-carboxylate (100.0 mg, 0.28 mmol, 1 equiv.) was dissolved in a mixture of ethanol, THF, and water (4:4:1), capped with a reflux condenser, and placed under an argon atmosphere. Iron powder (78.1 mg, 1.40 mmol, 5 equiv.) and ammonium chloride (37.4 mg, 0.70 mmol, 2.5 equiv.) were added to the reaction vessel at room temperature before the mixture was heated to 80 °C and stirred for at least 1 hour. Reaction progress was monitored with TLC (5% MeOH/DCM). Upon consumption of all the starting material, the reaction mixture was filtered hot, and the leftover residue was rinsed with a solution of 10% MeOH in DCM. The resulting filtrate was washed with water, dried over sodium sulphate ($Na_2SO_4$), filtered, and concentrated under reduced pressure. The product was purified using normal-phase chromatography on silica gel (0→20% MeOH/DCM) to provide a yellow-brown oil, which upon lyophilization, became pale yellow/ off-white powder (61.4 mg, 0.19 mmol, 67%).

$R_f$ = 0.52 (40% EToAc/DCM).

$^1$H NMR (400 MHz, DMSO-$d_6$) δ 8.48 (s, 1H), 7.47 (d, J = 8.8 Hz, 2H), 6.57 (m, 2H), 4.23 (q, J = 7.1 Hz, 2H), 1.24 (t, J = 7.1 Hz, 3H).

$^{13}$C NMR (100 MHz, DMSO-$d_6$) δ 163.12, 154.02, 129.58, 113.09, 60.99, 14.60.

LRMS m/z: $[M + H]^+$ calculated for $C_{12}H_{14}N_3O_4S_2^+$ 328.0420 ; Found 328.2000.

HRMS (ESI) m/z: $[M - H]^-$ calculated for $C_{12}H_{12}N_3O_4S_2^-$ 326.0275; Found 326.0283.

**5-((4-aminophenyl)sulfonamido)thiazole-4-carboxylic acid (MLLB-2201)**

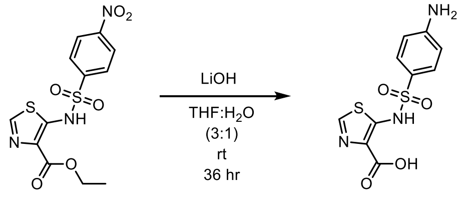


Ethyl 5-((4-aminophenyl)sulfonamido)thiazole-4-carboxylate (90 mg, 0.27 mmol, 1 equiv.) was dissolved in THF and water. To this, lithium hydroxide monohydrate (26.34 mg, 1.10 mmol, 4 equiv.) was added at room temperature and the mixture was left to stir for 36 hours. The mixture was diluted with water and washed with ethyl acetate. The aqueous layer was acidified with 1M hydrochloric acid to approximately pH 4, lest the free amine become protonated, and the product remain in the aqueous layer. Literature described the product precipitating out upon acidification, but this approach was found to be ineffective in readily isolating the target. Instead, the aqueous layer was then extracted with DCM and concentrated down to obtain an off-white solid. Should protonation of the free amine have occurred, the pH was adjusted using mild base (e.g., aqueous sodium bicarbonate). Location of the product was monitored using TLC. The obtained solid was dissolved in equal amounts of water/acetonitrile and purified using high pressure liquid chromatography (5 mM ammonium acetate: acetonitrile) with an Agilent 1290 Infinity II Preparative HPLC. Note that the product was found to be very heat sensitive and capable of undergoing decarboxylation quite readily. The collected HPLC fractions were freeze-dried to afford the product, MLLB-2201 as an off- white solid (12.3 mg, 0.04mmol, 15%)

$R_f$ = 0.## (5% MeOH/DCM).

$^1$H NMR (400 MHz, DMSO-$d_6$) δ 7.97 (s, 1H), 7.35 (d, J = 8.5 Hz, 2H), 6.49 (d, J = 8.6 Hz, 2H), 5.65 (s, 2H).

$^{13}C$ NMR (175 MHz, DMSO) δ 172.52, 163.60, 157.99, 151.88, 139.18, 129.86, 128.06, 127.02, 112.87.

HRMS (ESI) m/z: $[M + H]^+$ calculated for $C_{12}H_{10}N_3O_4S_2^+$ 300.0107; Found 300.####.

**$^{1}$H and $^{13}$C NMR Spectra**

Ethyl 5-((4-nitrophenyl)sulfonamido)thiazole-4-carboxylate (##) ($^{1}$H NMR; 400 MHz; DMSO-$d_6$)

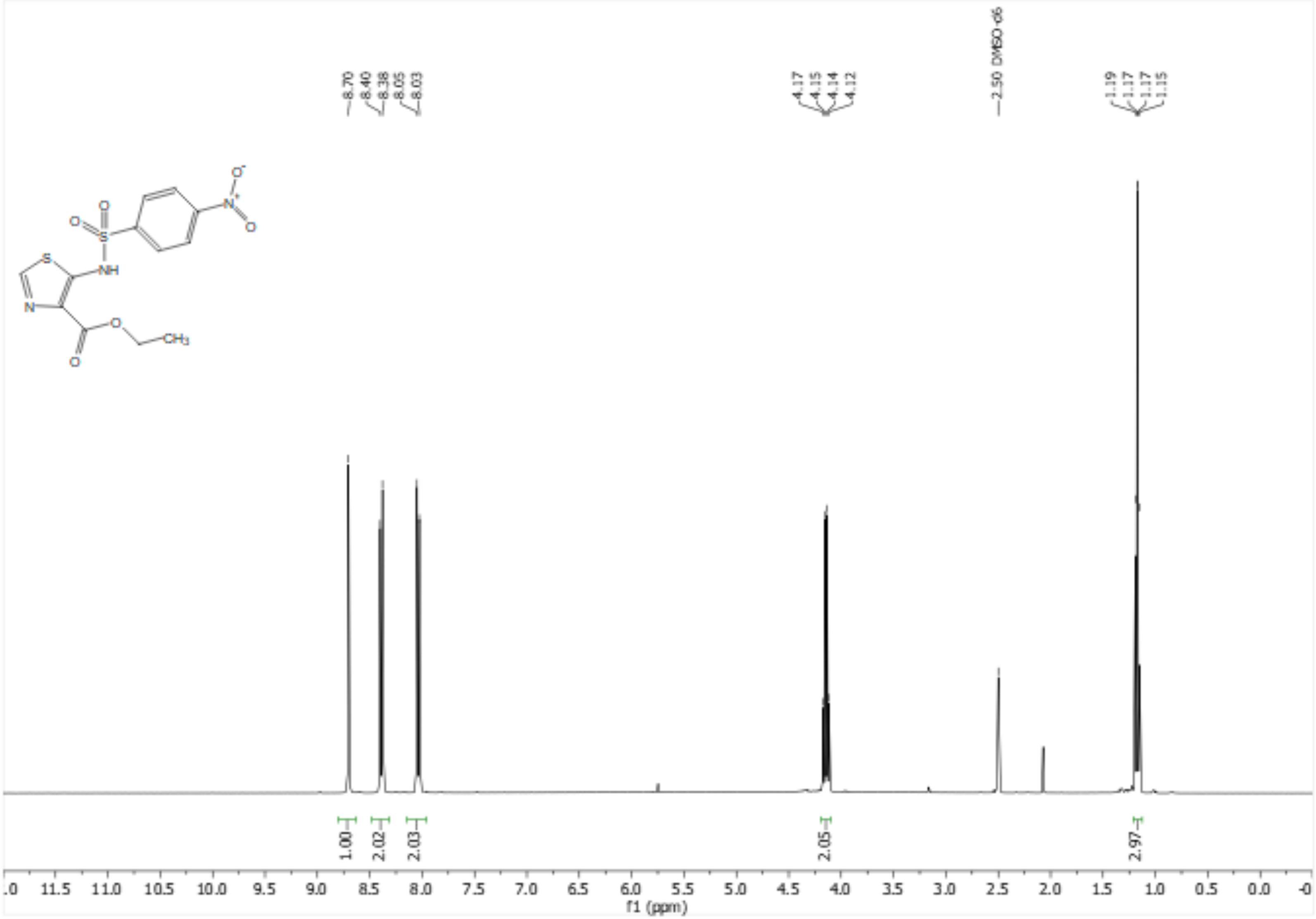

Ethyl 5-((4-nitrophenyl)sulfonamido)thiazole-4-carboxylate (##) ($^{13}$C NMR; 100 MHz; DMSO-$d_6$)

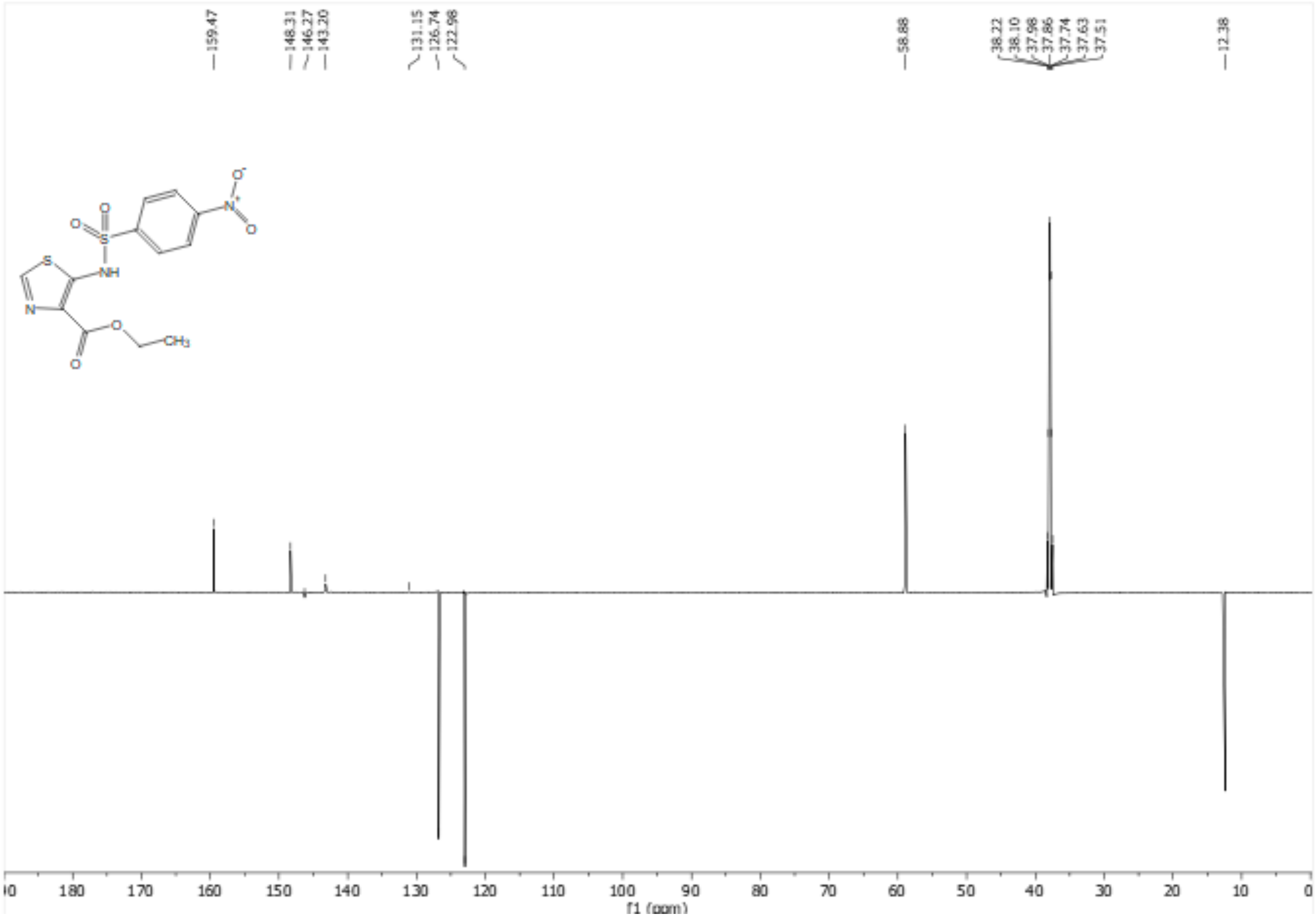

Ethyl 5-((4-aminophenyl)sulfonamido)thiazole-4-carboxylate (##) ($^1$H NMR; 400 MHz; DMSO-$d_6$)

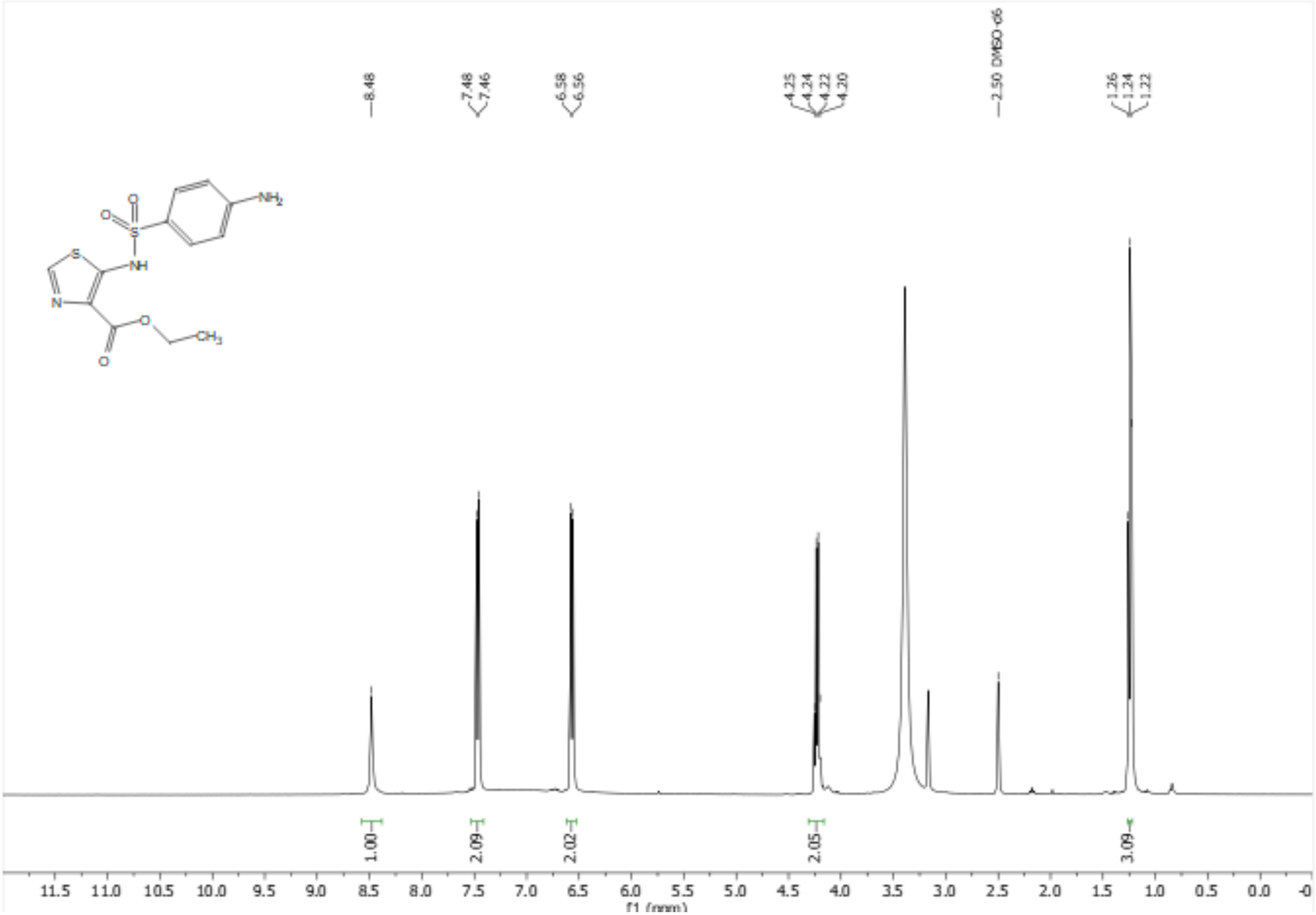

Ethyl 5-((4-aminophenyl)sulfonamido)thiazole-4-carboxylate (##) ($^{13}C$ NMR; 100 MHz; DMSO-$d_6$)

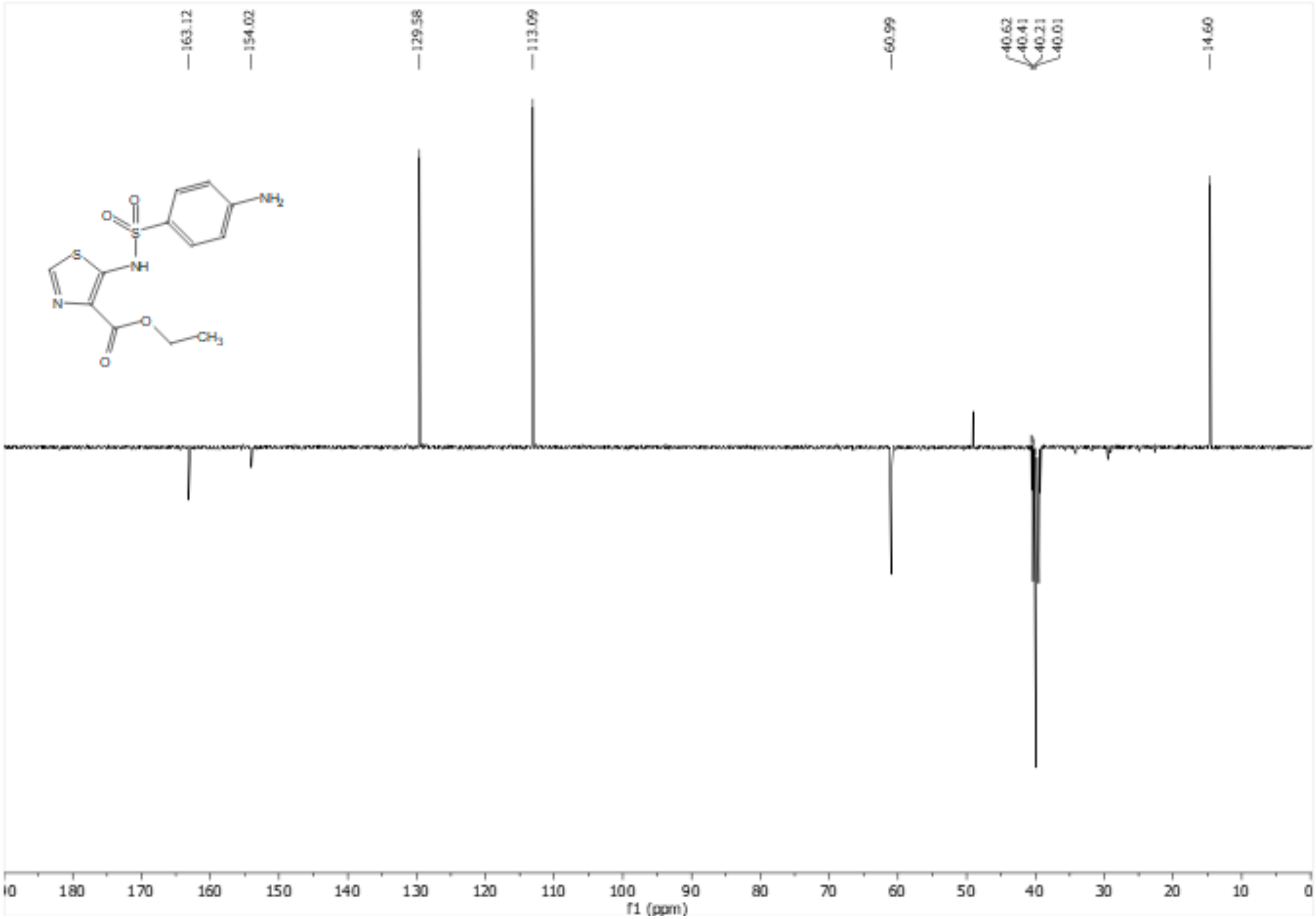

5-((4-aminophenyl)sulfonamido)thiazole-4-carboxylic acid (MLJB-2201) (##) ($^{1}$H NMR; 400 MHz; DMSO-$d_6$)

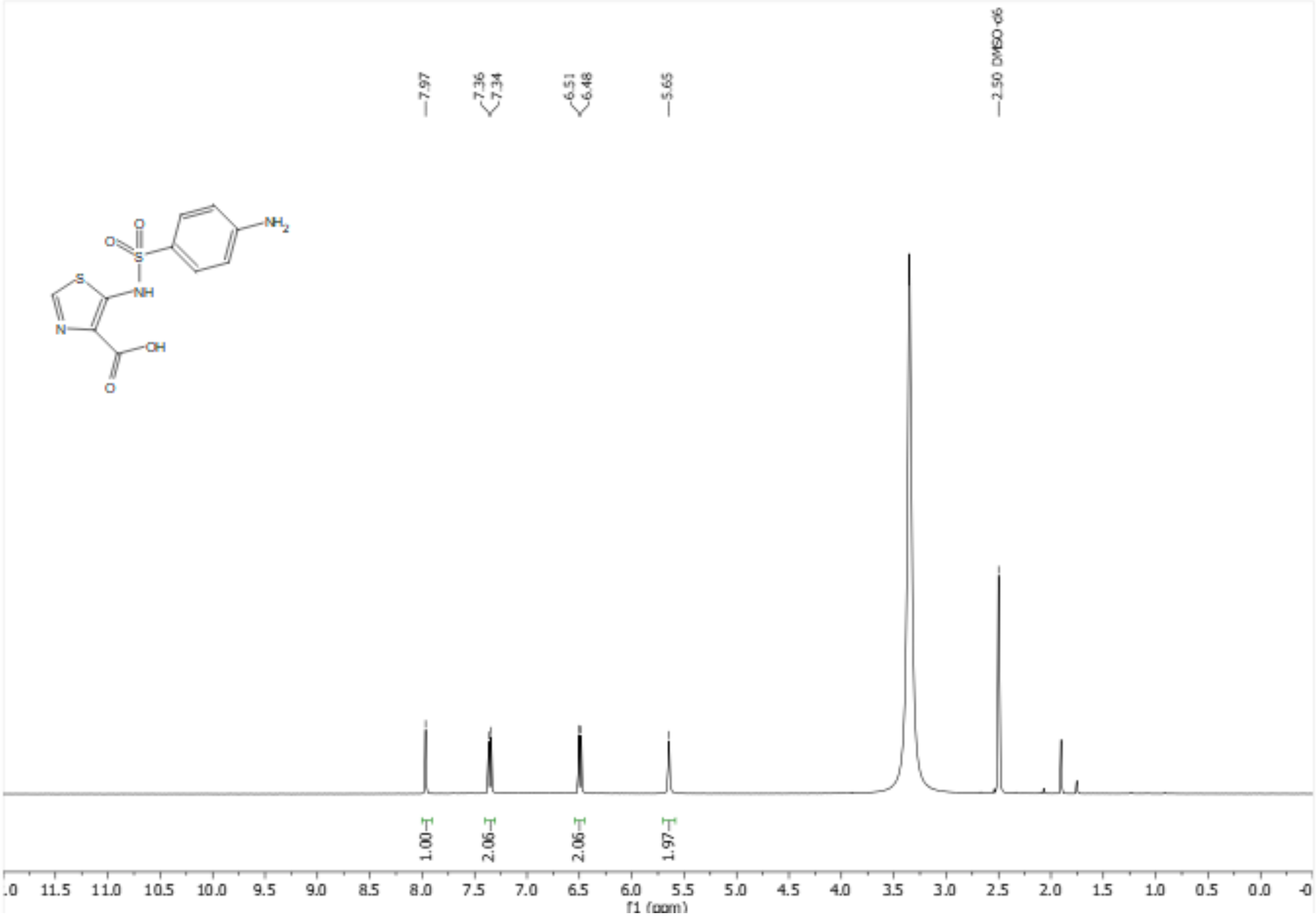

5-((4-aminophenyl)sulfonamido)thiazole-4-carboxylic acid (MLJB-2201) (##) ($^{13}C$ NMR; 100 MHz; DMSO-$d_6$)

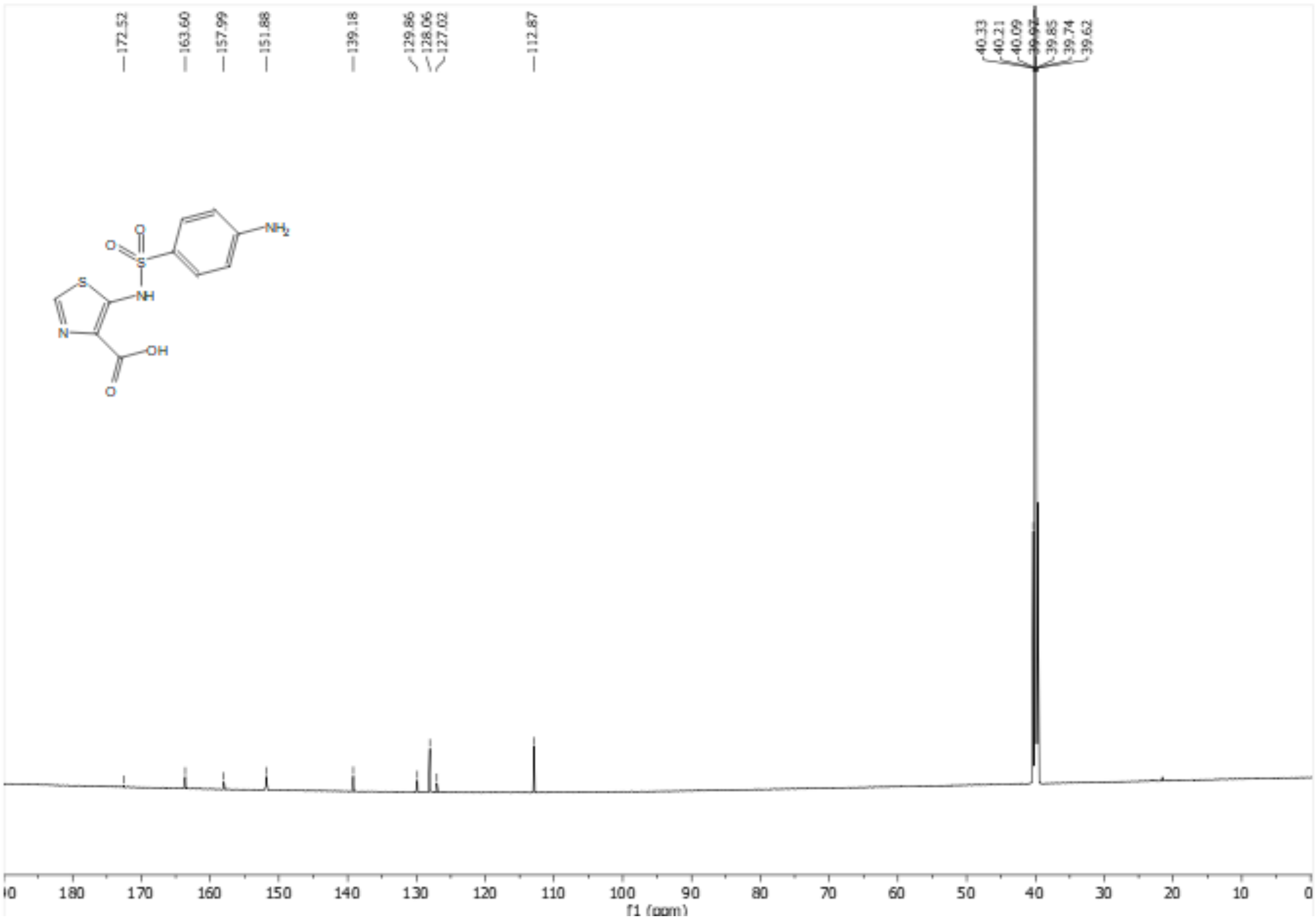
