## Supplementary material for "Metabolic connections between folate and peptidoglycan pathways in *Pseudomonas aeruginosa* inform rational design of a dual-action inhibitor": Yaeger et al. Supplementary Table S1

**Supplementary Table and References for Yaeger et al., Metabolic connections between folate and peptidoglycan pathways in *Pseudomonas aeruginosa* inform rational design of a dual-action inhibitor, Nov 22 2023**

**Table S1. Strains, plasmids, primers, and gBlocks**

| **Strains** | | |
|---|---|---|
| **Name** | **Description** | **Source** |
| PAO1 | Wild-type *P. aeruginosa* strain graciously donated by Keith Poole (Queen's University, Kingston, Canada) | [1] |
| *ampC* | An in-frame deletion mutant of *ampC* in PAO1 | This work |
| *ampR* | An in-frame deletion mutant of *ampR* in PAO1 | This work |
| *dacBC*** | A PAO1 mutant containing DacB S72A and DacC S64A catalytically inactive point mutations | This work |
| *dacBC**ampG* | A PAO1 mutant containing DacB S72A and DacC S64A point mutations and an in-frame deletion mutant of *ampG* | This work |
| *ampC* S90A | A PAO1 mutant containing an AmpC S90A catalytically inactive point mutation | This work |
| *oprF* | An *oprF* FRT mutant with the Gentamicin cassette flipped out | [2] |
| PAO1 +pUCP20 (Empty) | Wild-type PAO1 containing the pUCP20 empty plasmid. Carbenicillin resistant | This work |
| PAO1 +pUCP20 (*DHFRII*) | Wild-type PAO1 containing the pUCP20 plasmid expressing the *DHFRII* gene cloned from the pMS402 backbone. Carbenicillin resistant | This work |
| PA14 | Wild-type *P. aeruginosa* strain | [3] |
| *purC* | A PA14 mutant containing a transposon insertion in *purC*. Gentamicin resistant | [3] |
| *purD* | A PA14 mutant containing a transposon insertion in *purD*. Gentamicin resistant | [3] |
| *purF* | A PA14 mutant containing a transposon insertion in *purF*. Gentamicin resistant | [3] |
| *purH* | A PA14 mutant containing a transposon insertion in *purH*. Gentamicin resistant | [3] |
| *purL* | A PA14 mutant containing a transposon insertion in *purL*. Gentamicin resistant | [3] |
| *purM* | A PA14 mutant containing a transposon insertion in *purM*. Gentamicin resistant | [3] |
| *purN* | A PA14 mutant containing a transposon insertion in *purN*. Gentamicin resistant | [3] |
| *purT* | A PA14 mutant containing a transposon insertion in *purT*. Gentamicin resistant | [3] |
| *glpT* | A PA14 mutant containing a transposon insertion in *glpT*. Gentamicin resistant | [3] |
| *E. coli* K12 BW25113 | A wild-type *E. coli* strain with the genotype [F−Δ(*araD-araB*)567 *lacZ*4787Δ::*rrnB*-3 LAM− *rph*-1 Δ(*rhaD-rhaB*)568 *hsdR*514] | [4] |
| K12 + NDM1 | *E. coli* K12 BW25113 containing the pGDW plasmid expressing NDM-1. Kanamycin resistant | [5] |

| | | |
|---|---|---|
| *E. coli* DH5α | A wild-type *E. coli* strain with the genotype F– *endA1 gln*V44 *thi*-1 *recA1 relA1 gyrA96 deoR nupG purB20 φ80dlacZΔM15 Δ(lacZYA-argF)U169*, *hsdR17(rK–mK+)*, λ– | Invitrogen |
| *E. coli* SM10 | A wild-type *E. coli* strain used for efficient conjugative transfer of plasmid DNA to *P. aeruginosa* | [6] |
| *S. aureus* USA300 | A wild-type *S. aureus* strain. Methicillin resistant | [7] |
| **Plasmids** | | |
| **Name** | **Description** | **Source** |
| pMS402 (Empty) | A plasmid containing a promoter-less *luxBCDE* cassette. Kanamycin and trimethoprim resistant | [8] |
| pMS402Gm (Empty) | A plasmid containing a promoter-less *luxBCDE* cassette. Gentamicin and trimethoprim resistant | This work |
| pMS403 (Empty) | A plasmid containing a promoter-less *luxBCDE* cassette. Gentamicin resistant | This work |
| pMS403 (P*ampC*) | A plasmid containing the *luxBCDE* cassette under control of the *ampC* promoter. Gentamicin resistant | This work |
| pGDW (*NDM1*) | A plasmid expressing NDM-1. Kanamycin resistant | [5] |
| pUCP20 (Empty) | A plasmid with expression controlled by a *lac* promoter. Ampicillin/carbenicillin resistant | [9] |
| pUCP20 (*DHFRII*) | A plasmid with the *DHFRII* (*folA*) gene from pMS402 under control of a *lac* promoter. Ampicillin/carbenicillin resistant | This work |
| pEX18Gm (Empty) | A plasmid for performing allelic exchange. Gentamicin resistant | [10] |
| pEX18Gm (*ampC*) | The pEX18Gm plasmid containing the *ampC* deletion construct gblock. Gentamicin resistant | This work |
| pEX18Gm (*ampC* S90A) | The pEX18Gm plasmid containing the *ampC* S90A point mutation gblock. Gentamicin resistant | This work |
| pEX18Gm (*ampG*) | The pEX18Gm plasmid containing the *ampG* deletion construct gblock. Gentamicin resistant | This work |
| pEX18Gm (*ampR*) | The pEX18Gm plasmid containing the *ampR* deletion construct | This work |
| pUC57 (*ampC*) | The pUC57 plasmid containing the *ampC* deletion construct gblock. Ampicillin resistant | Genscript |
| pUC57 (*ampC* S90A) | The pUC57 plasmid containing the *ampC* S90A point mutation gblock. Ampicillin resistant | Genscript |
| pUC57 (*ampG*) | The pUC57 plasmid containing the *ampG* deletion construct gblock. Ampicillin resistant | Genscript |
| pPS856 | A plasmid containing a gentamicin resistance cassette within a multiple cloning site. Used to construct pMS402Gm | [10] |
| **Primers** | | |
| **Name** | **Sequence (5'-3')** | |
| P*ampC* Fwd | ATATGGATTCAGGGCGTTCAGCGGCAAATGG | |
| P*ampC* Rvs | TTAAGGATCCATTGGCGTCCTTTGTCGTTGGCTGC | |
| d*ampR* Up Fwd | TATAAAGCTTTCCGCCGTTTCGCCGCAATCTCC | |
| d*ampR* Up Rvs | GAAGCTTCGAAGGCGCGCAGGGCGTTCAGCGGCAAATGGGGTCGAACC | |

| | |
|---|---|
| d*ampR* Dwn Fwd | AACGCCCTGCGCGCCTTCGAAGCTTCCGGTTATGCAGGCGATTCAAGT GTCG |
| d*ampR* Dwn Rvs | AATTGGATCCGCTGGCTGGCGCGGTCGTCG |
| *DHFRII* Fwd | ATCAGAATTCGATTCACAAGAAGGATTCGACATGGG |
| *DHFRII* Rvs | ACTCAAGCTTCGTAAGATGCTTTTCTGTGACTGG |
| *purFtn Fwd* | ATCTTTCCAGCTTGAGCCAAGTAGGGG |
| *purFtn Rvs* | CCGCTCTCCCCTTTGTCAGTCG |
| *purDtn Fwd* | CGCCAGGAGAACCCCATGAACG |
| *purDtn Rvs* | AAGTCCCTTCGAAGGTGAATGGCCG |
| *purNtn Fwd* | CGTGGTCCTGAACAATCTGAAAAACC |
| *purNtn Rvs* | CATCTGTTTCCAGTCGGCGGTATAGC |
| *purTtn Fwd* | GGATTCCGTTTTCTTTCATTTCCGAGGTTCTTCC |
| *purTtn Rvs* | CGGCGCATCTCAGAGTTCGACG |
| *purLtn Fwd* | TTCCCGTCTCCAGAGGCTGTTCCG |
| *purLtn Rvs* | CGGCGAACCTCGGGAAGAAGGTCG |
| *purMtn Fwd* | GATCTCCGAATTACCCCTATAGGCCTGG |
| *purMtn Rvs* | TCAGCACCACGACATTGCAGAGTTTCG |
| *purCtn Fwd* | AGGCCCTATTACCCGTAAGCGGAGC |
| *purCtn Rvs* | TTCTTTCAGATGGTTGCGCGCGTTAGG |
| *purHtn Fwd* | CCAGCCCTCAGGACCCTTGC |
| *purHtn Rvs* | GCGTTATCCGCCCTGCGTGAAATTCG |
| **gBlocks** | |
| **Name** | **Sequence (5'-3')** |
| *ampC* deletion construct | gaattcTCAGCCAGTAGCTGCCGGTACTCACTTCGGTGGCGAACGGACGC CGGATGCTCTCGCTGGCCAGTTGCCGGGCAAACATCGCCGCCGGCGC CAGGGCCACGCCGACACCCTGGCGGGCCGCCTCGAGCATGGCCAGC GAGGTGTCGAAGACGATGCTCCGGGTCAGTGGCGCGTGCGCCGGCA GTCCGGCCGCCTGGAACCACAGCGGCCACTCGTCGGCGCGGTAGGA GCGCAGCAGGGTGTGCTGCAGCAGGTCGGCGGGACTGTGCAACTGG GCGGCGACCTCCGGGCAGCAGAGCACCGTCAGCGGCGCCTCGAACA ACGCCAGCGCCTCGGTGCCGTGCCAGGCGCCGCCGCCGAAGCGGAT CGCGTAGTCGAGCCCCTCGGCGGCGATGTCGACGCGGTTGTTGTGGG TGGACAGGCGCAGATCGATGAAGGGATGGCGCGCCTGGAAGTCCTCC AGCCGCGGCAGCAGCCAACCGACCGTGAAGGTTCCGACCGCGCCGA CGGTGAGCACGTCCCGGTAGTGGCCACCCTCGAAACGTTCCAGCAGG CCGGCGATGCGGTCGAAGGAGTCACACAGCACCGGCAGCAGGCTCTC GCCCTCGTGGGTCAGCATGAGGCCGCGCGGCAGACGCTTGAACAGG GCCACGCCGAGACGCTCCTCGAGGCTCTTCACCTGGTGGCTGACCGC CGCCTGGGTCACGCACAGCTCGATGGCCGCGCGGGTGAAGCTCAGGT GCCGGGCCGCTCGCGAGGGCGACGGAGCGTAGCGGCGCGGGACGC CGGTCCTGGCTATGATGGTGCCATGAGCGCTTCCCCGCCCCTCCGCCA GCCCTGCCCGCCCGGCGCCTGCGTCTGCGAACGCGAGCGCCTGGAG GCGCCCGGCGCGGACCGTCGCATCCTCCTCCTGACCCGCCAGGAAGA GCAGCGCCTGGCCGCTCGCCTGGAAGCCCTGCGCAGCCTGGAAGAC CTGGAACACCTGCTGCGGCGCATGGAGGAACAACTGGGCATCCGCCT GCGGATCGCCCCGGCCTTCGGCGAGGTGCGCAGCATGCGCGGCATCC GCATGCGCTTCGAGGAGCAGCCGGGGCTCTGCCGCAAGACCCGCCA GGCGATTCCCGCCGCCATCCGCCGCGGCCTGGAGAAGCGCCCGGAA GTCGCCTACGCCCTGCTCAACGCCCATGACCTGCTGCGCGACGCCTG |

| | |
|---|---|
| | AAGGTACTGAACGACAGGAAGAGGATGTCGCTCAAGACCTGGGCGGG CTTCAGGAGTATCGGCGGATAACGCCCATGGCGTTATTCGCCCTACAGG CCGCAGCGGATGCAGGCGAGCCCCGGGTCCGCCTGAATCCTTGGCGG GACTGGCCACCGCGCCGGGGTTCAGCTCGCCGACTCCGCCTGCACGT CGCCGAACATCGCCTGCAGGCGGAGCAGGCAGGCATCGCACAGGGTG CGCAGTTCGTCGAGGCGGATGCCGGCGAGGATTTCCATGCCGCGCAG CGGATCGGCGaagctt |
| *ampC* S90A point mutation | gaattcATGCGCGATACCAGATTCCCCTGCCTGTGCGGCATCGCCGCTTC CACACTGCTGTTCGCCACCACCCCGGCCATTGCCGGCGAGGCCCCGG CGGATCGCCTGAAGGCACTGGTCGACGCCGCCGTACAACCGGTGATG AAGGCCAATGACATTCCGGGCCTGGCCGTAGCCATCAGCCTGAAAGGA GAACCGCATTACTTCAGCTATGGGCTGGCCTCGAAAGAGGACGGCCGC CGGGTGACGCCGGAGACCCTGTTCGAGATCGGCGCGGTGAGCAAGAC CTTCACCGCCACCCTCGCCGGCTATGCCCTGACCCAGGACAAGATGCG CCTCGACGACCGCGCCAGCCAGCACTGGCCGGCACTGCAGGGCAGC CGCTTCGACGGCATCAGCCTGCTCGACCTCGCGACCTATACCGCCGGC GGCTTGCCGCTGCAGTTCCCCGACTCGGTGCAGAAGGACCAGGCACA GATCCGCGACTACTACCGCCAGTGGCAGCCGACCTACGCGCCGGGCA GCCAGCGCCTCTATTCCAACCCGAGCATCGGCCTGTTCGGCTATCTCG CCGCGCGCAGCCTGGGCCAGCCGTTCGAACGGCTCATGGAGCAGCAA GTGTTCCCGGCACTGGGCCTCGAACAGACCCACCTCGACGTGCCCGA GGCGGCGCTGGCGCAGTACGCCCAGGGCTATGGCAAGGACGACCGC CCGCTACGGGTCGGTCCCGGCCCGCTGGATGCCGAAGGCTACGGGGT GAAGACCAGCGCGGCCGACCTGCTGCGCTTCGTCGATGCCAACCTGC ATCCGGAGCGCCTGGACAGGCCCTGGGCGCAGGCGCTCGATGCCACC CATCGCGGTTACTACAAGGTCGGCGACATGACCCAGGGCCTGGGCTG GGAAGCCTACGACTGGCCGATCTCCCTGAAGCGCCTGCAGGCCGGCA ACTCGACGCCGATGGCGCTGCAACCGCACAGGATCGCCAGGCTGCCC GCGCCACAGGCGCTGGAGGGCCAGCGCCTGCTGAACAAGACCGGTT CCACCAACGGCTTCGGCGCCTACGTGGCGTTCGTCCCGGGCCGCGAC CTGGGCCTGGTGATCCTGGCCAACCGCAACTATCCCAATGCCGAGCGG GTGAAGATCGCCTACGCCATCCTCAGCGGCCTGGAGCAGCAGGGCAA GGTGCCGCTGAAGCGCTGAaagctt |
| *ampG* deletion construct | gagctcGTTGCAGTTCACCAACTTCCGCGATTGACGAGGTTTTTCGGAAC CTCTCGCCATCCCGGGCATCGCAATAGGGAGTAGCGCCGCCGCGACC TCTCCCGCGTTGTCCGCCCCTGTCGGCGCAGCGTCGGTTCGTCCGCG CTGCGCCGATTCGTATCCAGGCCCGGAAACGTCGCCCCACGGCGTCC GGGCAGGAATCTGCAAGCCCGTCCCGTTTCCGTTGCGGAAGGGAGGG GTGTTTTTTCGAGAAATGGAGGTAATGCATGGAATTGAACTACGACCGA CTGGTGCAGCAGACCGAGTCCTGGCTGCCGATCGTGCTGGAGTACAG CGGCAAGGTCGCCCTGGCGCTGCTGACCCTGGCGATCGGCTGGTGG CTGATCAACACCCTGACCGGCCGGGTCGGCGGCCTGCTCGCCAGGCG CAGCGTCGACCGCACCCTGCAAGGCTTCGTCGGCAGCCTGGTGAGCA TCGTCCTGAAGATCCTGCTGGTGGTCAGCGTGGCTTCCATGATCGGCA TCCAGACCACCAGCTTCGTCGCCGCCATCGGCGCCGCCGGCCTGGCC ATCGGCCTGGCCCTGCAGGGCAGCCTGGCTAACTTCGCCGGCGGCGT GCTGATCCTGCTGTTCCGCCCGTTCAAGGTCGGCGACTGGATCGAGG CACAGGGCGTGGCCGGCACCGTGGATTCGATCCTGATCTTCCACACCG TGCTGCGTAGCGGCGACAACAAGCGGATCATCGTGCCCAACGGGGCG CTGTCCAACGGAACGGTGACCAACTACTCCGCCGAGCCGCTGCGCAA GGTGGTCTTCGACGTCGGCATCGACTACGACGCCGATCTGAAGAATGC |

| | |
|---|---|
| | GCAGAACATTCTCCTGGCCATGGCCGACGATCCGCGGGTTCTGAAGGA<br>CCCGGCACCGGTGGCGGTGGTTTCCAATCTCGGCGAAAGCGCGATTA<br>CCCTGTCCCTGCGGGTCTGGGTGAAGAACGCCGACTACTGGGACGTG<br>ATGTTCATGTTCAACGAAAAGGCCCGCGACGCGCTGGGCAAGGAAGGT<br>ATCGGCATTCCCTTCCCGCAGCGGGTGGTCAAGGTTGTGCAGGGCGC<br>GATGGCCGACTGAGGTCCGTGCGATCCACGAAAAAGGCCGGCGAATG<br>CCGGCCTTTTTCATTCTCGCCTCTAGACGCAAAAAATAACGCGCACTCT<br>AACCGCTCTACTTCGCTGTAAGCCAGCCATGCCGACGTATGTCCACCCT<br>GTCGTTGTGCAGGAGAACGCCTTCCTCGCGCAAGCGTGCGCGTTGTT<br>CGCGACCGCCGGGACTGTCCGCCGGCAGGCTGATGCGACCGCCGGC<br>GCCGAGAACCCGATGCCAGGGCAGGCGGGTATCCACCGGCAACTGGC<br>TGAGCGTCCTGCCGACCCAGCGGGCGGCGCGGCCTAGGCCGGCCAG<br>TTCGGCCAGTTGCCCGTAGCTGACCACCTGCCCCGGCGGCACCTGCG<br>CCAGCACCAGGTACAGGGCCTCGCGGCGCGCCTGGGCACTTGCCGG<br>GTCGCTGCCGGCCCATGCATGCTCATCGGTTTTCCCTTTCCTGTCCGC<br>CTTGTGAACGCCCATGTCGCTCAGAGCCCTCTGTTCAATCGTCGTGAT<br>GCTCCTGGCCGGCCCCGCGCTGGCCGACACCGTATGGCTGGACAACG<br>GCGACCGCCTGTCCGGCGAGATCGTGCTGATGGACGGCGGCAAGCTG<br>GCGCTGAAGACCCGTTATGCCGGCCAGGTGCTGATCGATTGGAAGGAT<br>ATCGACACCATCAGTTCCGACAAACCCCTGCTGATCAAGCAGCAGGGC<br>GTATCCGGGCAGCGTAGCCGAACCCTGGAGGCGGCGGGCAAGGGCAT<br>GGTGCGGATCGTCGATGGCGGCAGCCATACCGTCCCGCTGGCCAGTAT<br>TCGCCAGATGGTGCCGCCGCGACCGCTGGTGGAGGACCTGGTCTGG<br>GAGGGCAATCTCGACGTCAAGCTGGATAGCAAGCGCAACGACAGCGA<br>CAAGGACGAATGGAAGCTCAAGGGCGATACCCGGCTTCGCCACGGTG<br>CCTGGCGCCACGTGCTGGCGGGGGAAGTGGAGCGGGAAAAGAAGGA<br>CGGGCGCAAGGTCGAGGACAACTGGGAGCTGGACTACGACCTCGATC<br>GCTTCTTCGACGAACACTGGTTCTGGCGCGGCAGCTATTCGCAGAAGC<br>ACGATGCGATCGACAACCTGGAGCGCCAGAGCGCGCTGGGAACCGGC<br>CCCGGCTACCAGTTCTGGGACGACGAACTCGGGCGTTTCGACCTGGT<br>CGCCGAGATCAGTCGCTGGCAACTGGAGTGGCGaagctt |
